# Stressed β-cells contribute to loss of peri-islet extracellular matrix in type 1 diabetes

**DOI:** 10.1101/2025.06.25.661601

**Authors:** Chelsea G. Johansen, Kenedee Lam, Nikki L. Farnsworth

**Affiliations:** Department of Chemical and Biological Engineering, Colorado School of Mines, Golden, CO; Quantitative Biosciences & Engineering, Colorado School of Mines, Golden, CO

## Abstract

Type 1 diabetes (T1D) is characterized by the immune-mediated destruction of insulin-producing β-cells in pancreatic islets. The peri-islet extracellular matrix (ECM) is a complex protein barrier that is lost in T1D, in part due to infiltrating immune cells. The contribution of stressed β-cells to ECM degradation during T1D remains unclear. To bridge this gap, we used 12–15-week-old NOD mice and pancreas sections from healthy, ≥2 autoantibody positive (Aab+), and recent onset T1D donors. We focused on MMP-3 due to its role in degrading type IV collagen (COL IV) in the peri-islet ECM. Treatment with proinflammatory cytokines or hyperglycemia increased MMP-3 gene expression and protein levels in mouse and human islets. In NOD pancreas sections, increased MMP-3 expression in β-cells correlates with loss of COL IV during insulitis and hyperglycemia; however, this was independent of insulitis score. We observed similar increases in MMP-3 and loss of COL IV in islets and exocrine tissue from Aab+ and recent onset T1D donors. These results suggest that stressed β-cells degrade the ECM during preclinical T1D, further weakening the peri-islet ECM barrier and facilitating islet infiltration and death. Inhibiting expression of MMP-3 may represent a novel treatment to prevent islet death in T1D.

## Introduction

The peri-islet extracellular matrix (ECM) provides structural and biochemical support to pancreatic islets that maintains cellular function and homeostasis^1,2^. Islets are surrounded by two forms of specialized ECM: the basement membrane (BM) that forms a capsule directly around the islet and the interstitial matrix (IM) that connects the exocrine and endocrine compartments around peri-islet BM, major ducts, and arteries^3–5^. BM is primarily composed of laminin-10, type IV collagen (COL IV), fibronectin, and IM is composed of fibrillar collagens and highly sulfated proteoglycans^3,6–8^. Together, the BM and IM provide biochemical and mechanical cues that promote islet viability and function^6,9–11^.

Type 1 diabetes (T1D) is an autoimmune disease characterized by the immune-mediated destruction of insulin-producing β-cells, leading to chronic hyperglycemia^12,13^. The disease progresses through defined stages, beginning with the presence of islet-directed autoantibodies (stage 1), followed by dysglycemia due to impaired insulin secretion (stage 2), and ultimately clinical diagnosis (stage 3), when 60%–90% of β-cell mass has been lost^12,14^. While the immune-mediated attack on β-cells has been extensively studied within the context of T1D, emerging evidence suggests that loss of the peri-islet ECM plays a crucial role in disease onset and progression. Degradation of the ECM is mediated by proteolytic enzymes, like matrix metalloproteinases (MMPs), A Disintegrin and Metalloproteinase with Thrombospondin Motifs (ADAMTS) proteases, and heparanases^5,15–19^. These enzymes play crucial roles in tissue repair and disease progression by degrading structural ECM components, such as COL IV and laminin-10 as found in the peri-islet BM, which allows for immune infiltration and progression of T1D^3,4,6,20^. Activated immune cells have been implicated in the secretion of heparanases and gelatinases, which degrade fibrillar collagens (type I and type III), elastin, and fibronectin^20–25^. Previous work has shown that treating prediabetic NOD mice with a heparanase inhibitor, aimed at preventing heparan sulfate degradation in the peri-islet BM, led to preservation of heparan sulfate within the islet and protected islets from destructive autoimmunity and T1D^5^. This strongly suggests that loss of the peri-islet ECM is a critical step in the pathogenesis of T1D.

Although changes to the peri-islet ECM have been well characterized in NOD mice and human donors with T1D 2+ autoantibodies, how these changes contribute to T1D pathogenesis, particularly the sharp increase in β-cell death from stage 1 to stage 3 of T1D, remains poorly understood. Previous studies have characterized loss of ECM proteins, including COL IV, laminin, nidogen, and perlecan in NOD mice and human donors with 2 or more Aab+ as well as longstanding T1D^3,4^. Moreover, ECM loss during immune infiltration has been attributed to matrix-degrading enzymes secreted by immune cells^3^. However, β-cells have also been implicated in expressing these enzymes upon chemical stress and interactions with immune cells^24^, though their role in matrix degradation in T1D-associated stress has yet to be investigated. While it has been observed that islets with higher levels of immune cell infiltration have greater loss of peri-islet ECM, this does not fully explain the global loss of ECM components in both the exocrine and endocrine compartments of the pancreas in T1D^15–20^. The observed loss of ECM proteins leads to the elimination of critical signaling pathways associated with islet function and survival^3,6^. Therefore it is critical to understand the cells responsible for degrading the ECM in T1D to preserve islet function and potentially prevent disease progression. Additionally during islet infiltration, T-cells and macrophages produce high levels of pro-inflammatory cytokines, including interferon-α (IFN-α), interleukin-1β (IL-1β), and interferon-γ (IFN-γ), that cause islet stress, dysfunction, and death^26,27^. Cytokine induced cellular stress causes upregulation of tissue degrading proteases in a number of cell types, but cytokine-induced ECM degradation by the β-cell has not been explored^28^. Together, these findings suggest that peri-islet BM degradation contributes to β-cell destruction and that preserving BM could be a potential therapeutic strategy for preventing T1D.

It is currently unknown if β-cells have a role in degrading the peri-islet ECM in T1D and if they contribute to the degradation of their ECM. This study aims to investigate the temporal aspects of ECM loss during T1D pathogenesis and the role of stressed β-cells in actively degrading their ECM. The focus of our study is MMP-3, a broad substrate stromelysin-1 that degrades COL IV and is known to activate other MMPs, thus amplifying ECM remodeling in inflammatory conditions^15,18,28,29^. We investigated the contributions of β-cell stress driven by hyperglycemia, insulitis, and cytokine exposure, to the degradation of COL IV in the peri-islet ECM during T1D. Determining if β-cells actively remodel their microenvironment will provide evidence that they may contribute to their own decline in T1D and that this may contribute to the sharp decline in β-cell mass that is observed prior to clinical diagnosis of T1D. A deeper understanding of this process could open new avenues for therapeutic intervention, potentially slowing or halting disease progression, preserving remaining β-cells, and creating conditions that support their regeneration.

## Results

### Proinflammatory Cytokines and Hyperglycemia Lead to Upregulation of ECM Degrading Enzyme MMP-3 in Mouse and Human Islets

To understand how cytokine stress in T1D affects islet function, we used qPCR to analyze gene expression of MMP-3, the broad substrate stromelysin which degrades COL IV^30^. Mouse and human islets were treated with and without a 0.1X proinflammatory cytokine cocktail (1ng/ml TNF-α, 0.5ng/ml IL-1β, 10ng/ml IFN-γ) for up to 72h. This concentration of cytokine cocktail has been shown to cause dysfunction to insulin secretion without significant impacts on islet viability in previous studies^31^. In cytokine-treated mouse islets, there is a 23.92 ± 3.85 and 33.04 ± 6.89-fold increase in MMP-3 gene expression at 48h and 72h, repsectively, when compared to the control (p=0.027, p=0.043, Figure 1A). In cytokine-treated human islets, there is a 167.7 ± 45.08 and 208.7 ± 23.66-fold increase in MMP-3 gene expression at 48h and 72h, repsectively, when compared to the control (p=0.066, p=0.013, Figure 1B). These findings support that proinflammatory cytokine stress upregulates MMP-3 gene expression in both mouse and human islets. To further demonstrate that MMP-3 expression is elevated in islets with T1D, 12-15-week-old normoglycemic NOD mouse islets and age matched NOD-RAG1KO control mouse islets were isolated for qPCR. Our data demonstrates a 4.57 ± 1.99-fold increase in MMP-3 gene expression in normoglycemic NOD mice normalized to the immunodeficient NOG-RAG control (Figure 1C).

**Figure 1:**
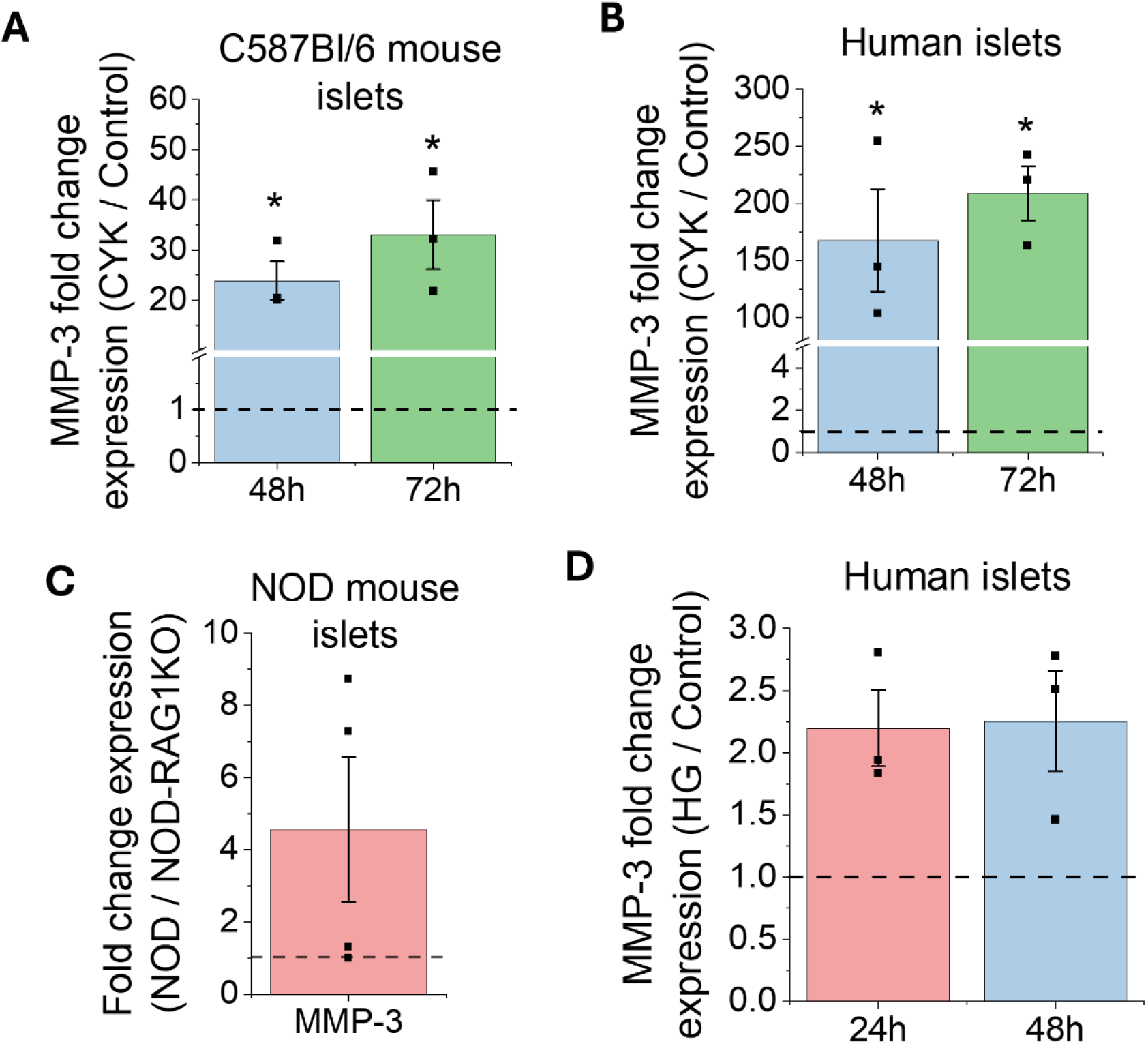
Cytokine treatment upregulates MMP-3 gene expression in both mouse and human islets. (A) MMP-3 fold change between 0.1X cytokine (CYK) treated mouse islets and untreated mouse islets (n=3). (B) MMP-3 fold change between 0.1X CYK treated human islets and untreated human islets (n=3). (C) MMP-3 fold change between normoglycemic NOD mouse islets and immunodeficient, age-matched NOD-RAG1KO mouse islets (n=4). (D) MMP-3 fold change between hyperglycemic human islets and untreated control human islets at 24h and 48h (n=3). The ΔΔCt method was used to calculate fold change between treatment groups. HPRT1 is the housekeeping gene for all qPCR experiments. Error bars represent the mean +/- SEM. * indicates statistical significance as determine by the 95% confidence interval.

To understand how hyperglycemia, associated with the sympotmatic phase of T1D, affects islet function, human islets were cultured with excess 20mM glucose for up to 72h to induce hypeglycemia. Glucose-stimulated insulin secretion (GSIS) was used to determine hyperglycemia induced islet stress (Figure S1A-B). The presented GSIS data is a measure of insulin secreted to the bulk media normalized to insulin content in the islets. In the 72h hyperglycemic (HG) condition, there is an increase in insulin secreted at 2mM glucose and a decrease in insulin secretion at 20mM glucose compared to untreated controls, where this dysfunction was most notable at 24h for the 20mM glucose treatment and at 72h for the 2mM glucose treatment, indicating dysfunction to GSIS (Figure S1A). We also calculated the stimulation index (SI), which is the ratio of insulin secreted at 20 mM versus 2 mM glucose, to assess β-cell responsiveness to glucose. The SI decreased from the control group to the HG group across all time points (p<0.0005, Figure S1B). Specifically, at 72h, the SI in the HG condition was significantly lower than the 72h control (p=0.0053, Figure S1B), which further demonstrates dysfunction to insulin secretion. Our GSIS findings are supported by previous studies showing that islets cultured in excess (16-28 mM) glucose exhibit increased insulin release at low (2-4 mM) glucose levels and fail to further increase insulin release when exposed to high (16.7 mM) glucose^32,33^. These results confirm that culturing islets in high glucose is a useful model for mimicking the effects of chronic hyperglycemia on human β-cell function. Having established β-cell dysfunction under hyperglycemia, we next examined how this stress affects MMP-3 gene expression. The qPCR data demonstrates a 2.20 ± 0.31 fold increase at 24h and a 2.25 ± 0.40-fold increase at 48h in MMP-3 gene expression in HG treated islets compared to control (Figure 1D). Together, the qPCR data supports a role for cytokine treatment and hyperglycemia in mediating the upregulation of MMP-3 in mouse and human islets.

We next assessed MMP-3 protein levels to confirm whether transcriptional upregulation resulted in increased protein expression. Mouse and human islets were treated with and without a 0.1X proinflammatory cytokine cocktail for up to 72h as described above. At 48 and 72h, MMP-3 protein levels increased by 1.48 ± 0.05 and 1.22 ± 0.04 fold repsectively in cytokine treated mouse islets compared to untreated controls (p=0.048, p=0.042, Figure 2A-B). Similar results were found in human islets, where we observed a 1.02 ± 0.33, and 2.02 ± 0.86-fold increase in MMP-3 protein expression at 48h and 72h between cytokine-treated and control conditions, respectively (Figure 2C-D). When human islets were treated with excess 20mM glucose in a hyperglycemic condition, there was a 1.17 ± 0.17 and 1.25 ± 0.08-fold increase in MMP-3 protein expression at 48h and 72h between hyperglycemic and control conditions, respectively (Figure 2E-F). Together, this data supports that proinflammatory cytokine stress and hyperglycemic conditions can lead to islet-specific MMP-3 protein expression in mouse and human islets.

**Figure 2:**
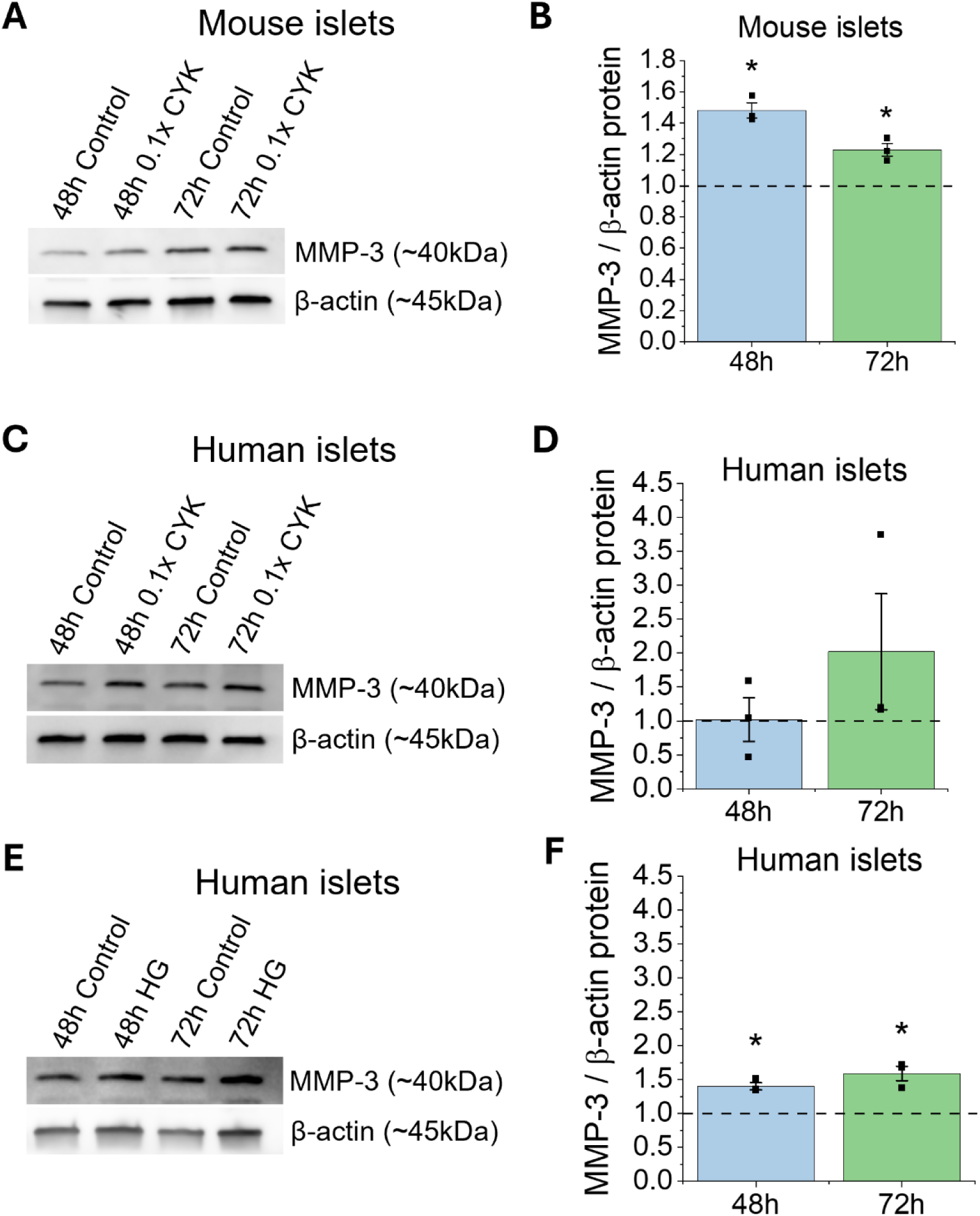
Cytokine treatment and hyperglycemia lead to islet-specific MMP-3 protein expression in mouse and human islets. (A) Representative western blot of MMP-3 and housekeeping protein β-actin in whole mouse islets untreated or treated with 0.1X cytokine (CYK) cocktail (n=3). (B) Western blot quantification for MMP-3 expression in mouse islets. (C) Representative western blot of MMP-3 and β-actin in whole human islets untreated or treated with 0.1X cytokine (CYK) cocktail (n=3). (D) Western blot quantification for MMP-3 expression in human islets. (E) Representative western blot of MMP-3 and β-actin in hyperglycemic (HG) or control human islets (n=3). (F) Western blot quantification for MMP-3 expression in human islets. For all western blot experiments, MMP-3 expression was normalized to β-actin expression and BCA assay total protein results. Error bars represent the mean +/- SEM. * indicates statistical significance as determine by the 95% confidence interval.

### Hyperglycemia-Induced Expression of MMP-3 and Degradation of COL IV in Mouse Pancreas Sections

After establishing that stressed islets express MMP-3 at the gene and protein level *in vitro*, we next sought to evaluate expression of MMP-3 in the NOD mouse, which spontaneously develops T1D after 12 weeks of age and exhibits insulitis starting from as early as 4 weeks of age. We examined MMP-3 and COL IV in pancreas sections from 12-15-week-old normoglycemic (NG) NOD mice, age matched hyperglycemic (HG) NOD mice, and age matched immunodeficient NOD-Scid mice using immunohistochemical staining of fresh frozen pancreas sections. MMP-3 and COL IV were stained in subsequent sections. Dense populations of cyan nuclear staining on the islet periphery were used to assign insulitis scores in combination with insulin staining (Figure 3A-B). The 12-15-week normoglycemic NOD mice had heterogeneous levels of insulitis both within a single animal as well as between animals. In the NG NOD and HG NOD, we see an expected decrease in insulin-positive area and an increase in immune infiltration indicating the onset of T1D in these animals (Figure 3A). Furthermore, there is evident MMP-3 expression that is increased in islets from NG NOD and HG NOD sections compared to NOD-Scid controls evidenced in the areas pointed out by white arrows in Figure 3A. We also noticed increases in exocrine MMP-3 expression in NG NOD and HG NOD compared to NOD-Scid controls; however islet staining for MMP3 appeared stronger than in the exocrine tissue (Figure 3A). In contrast, we see a decrease in COL IV staining in NG NOD and HG NOD sections compared to NOD-Scid controls, where areas of high immune infiltration have a lack of COL IV staining as indicated by white arrows in Figure 3B. Specifically, in the HG NOD, there is a decrease in COL IV in the insulin positive islet area as well as the infiltrated islet periphery. The COL IV within the vasculature appeared intact in NG and HG NOD mice; however, staining intensity is reducd suggesting that there is less BM surrounding vasculature compared to NOD-Scid controls (Figure 3B).

**Figure 3:**
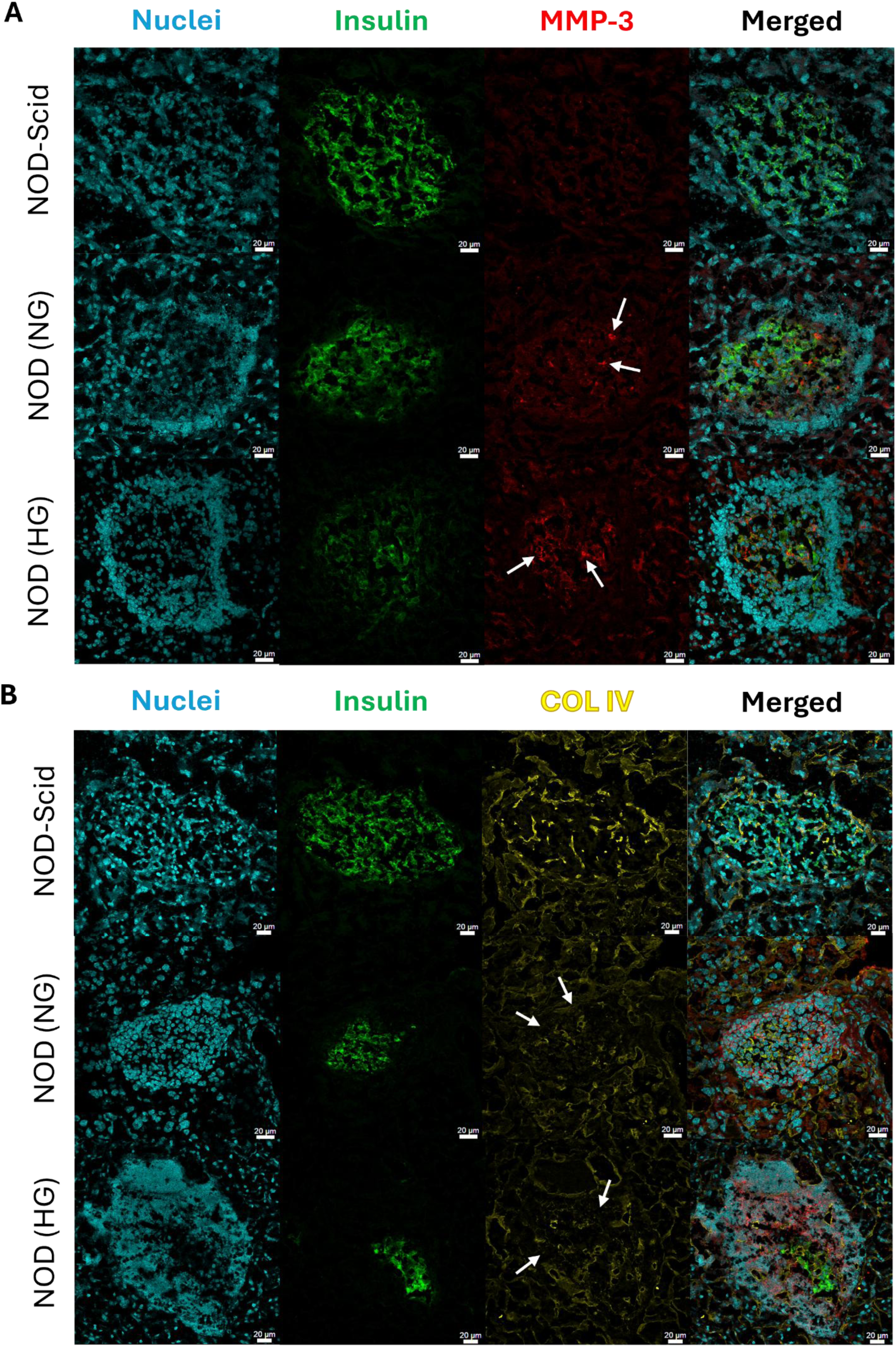
MMP-3 and COL IV IHC analysis in T1D mouse pancreas sections. Representative confocal microscopy images of (A) MMP-3 and (B) COL IV in immunodeficient NOD-Scid, normoglycemic (NG) NOD, and hyperglycemic (HG) NOD mouse pancreas sections. White arrows in A are pointing to positive MMP-3 staining within the islet and the white arrows in B are pointing to areas without COL IV. MMP-3 and COL IV staining was done on consecutive sections of tissue in each condition. Insulin staining in green was used to identify pancreatic islets. All scale bars are 20μm.

We next wanted to quantify changes in MMP-3 and COL IV in the tissue sections from Figure 3 to further support our observations. We calculated the total MMP-3 staining as the MMP-3 fluorescence intensity multiplied by staining area, normalized to insulin positive area, and found an increase in MMP-3 staining in both NG and HG NOD mice compared to NOD-Scid controls, where HG NOD mice had the highest levels of β-cell specific MMP-3 expression (p<0.0001, p<0.0001, Figure 4A). Stratification of MMP-3 levels by insulitis score yielded an increase in β-cell MMP-3 levels with increasing insulitis score for both the NG and HG NOD (p=0.034, Figure 4B). However, β-cell MMP-3 levels were consistenly higher in HG NOD sections compared to NG NOD sections and this effect was independent of insultis score (Figure 4B). Overall, our data suggests that the expression of MMP-3 in insulin-producing β-cells is driven by insulitis and to a larger degree hyperglycemic stress in the pre-diabetic NOD mouse.

**Figure 4:**
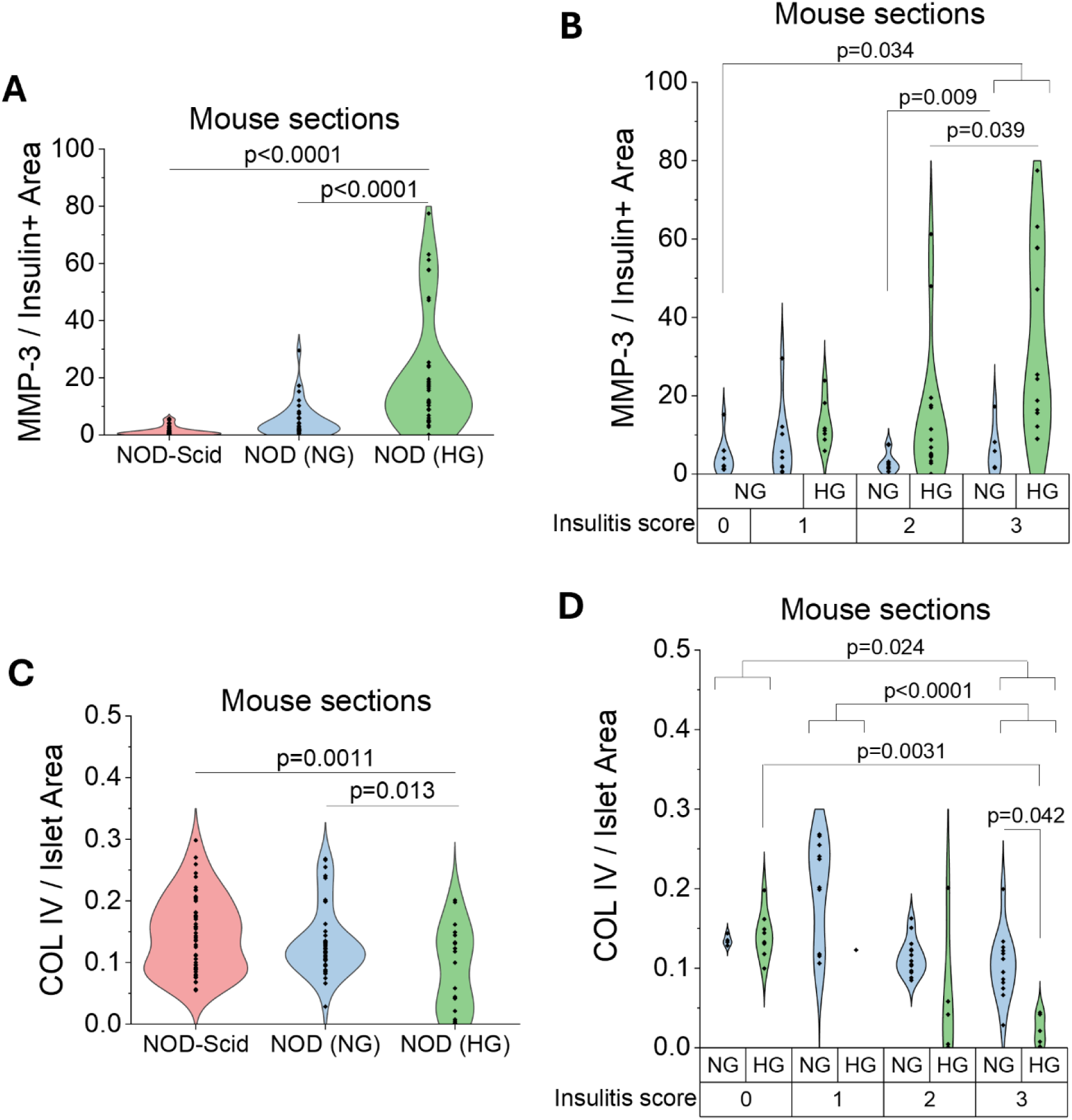
Hyperglycemia and insulitis-induced expression of MMP-3 and degradation of COL IV in mouse pancreas sections. (A) Quantification of MMP-3 intensity multiplied by staining area normalized to insulin positive area in the confocal images from Figure 3A for NOD-Scid, NG NOD, and HG NOD mouse sections. (B) NG NOD and HG NOD data in A stratified by insulitis score. (C) Quantification of COL IV-stained area normalized to total islet area in the confocal images from Figure 3B for NOD-Scid, NG NOD, and HG NOD mouse sections. (D) NG NOD and HG NOD data in C stratified by insulitis score. p<0.05 indicates statistical significance as determine by ANOVA with Tukey’s post hoc analysis.

We next quantified COL IV staining as the total COL IV positive area normalized to total islet area incluidng any immune infiltrates. We found a slight decrease in COL IV-stained area normalized to total islet area in the NG NOD sections compared to the NOD-Scid controls and a significant decrease in COL IV staining in the HG NOD (p=0.0011, Figure 4C). Stratification by insulitis score further revealed a progressive decline in COL IV-stained area from low (score 0–1) to high (score 3) infiltration across both normoglycemic and hyperglycemic conditions. While at low insulitis scores (0-1)mthere were not significant differences in COL IV area, likely due to low numbers of islets with this score in the HG NOD sections, in islets with high insulitis scores (2-3) the HG NOD sections had significantly less COL IV than NG NOD mice with the same insulitis score (Figure 4C). Together with the MMP-3 results, this supports that hyperglycemic stress upregulates β-cell MMP-3 expression leading to a loss of peri-islet COL IV that is independent of insulitis.

### Hyperglycemia-Induced Expression of MMP-3 and Degradation of COL IV in Human Pancreas Sections

To determine if human islets are also capable of remodeling the ECM in T1D, MMP-3 and COL IV were stained in human pancreas sections from healthy, two or more autoantibody-positive (Aab+), and recent onset (<6 months) T1D donors to determine whether similar trends in MMP-3 expression and COL IV degradation are conserved across species. Human donors were age-, sex-, and BMI-matched as closely as possible (Table 3). T1D donors were within six months of diagnosis and were likely hyperglycemic based on HbA1c levels ≥10. nPOD pathological analysis confirmed the presence of insulitis in Aab+ and T1D donor tissue blocks where sections were requested. Similar to our observations in NOD mice, we saw an increase in MMP-3 expression in human islets from Aab+ and T1D donors, where MMP-3 signal was also increased in the exocrine tissue (Figure 5A). In both Aab+ and T1D donors we also observed a decrease in COL IV staining both in the islet and exocrine tissue, where loss of the peri-islet BM was consistently observed in all samples (Figure 5B). In islets from T1D donors, we observed an increase in COL IV specifically in the islet vasculature (white arrows, Figure 5B); however, peri-islet and exocrine COL IV also slightly recovered around insulin positive islets in T1D donors (Figure 5C). Quantification of the images as described above confirmed that islet MMP-3 intensity multiplied by staining area normalized to insulin positive area was increased in Aab and T1D donors compared to healthy controls, where MMP-3 levels were highest in T1D donors that were presumed hyperglycemic (p<0.001, p<0.001, Figure 6A). The opposite trend is seen for COL IV-stained area normalized to total islet area where there is a decrease in COL IV in Aab+ donors (p<0.001) and T1D donors (p=0.017, Figure 6B) compared to healthy controls. Further, we see a sligtht increase increase in COL IV levels in the islet in T1D donors compared to Aab+ donors (p=0.017, Figure 6B) that corresponds to the increase in vascular COL IV observed within the islet and is not associated with recovery of the peri-islet BM (Figure 5B). We also quantified exocrine COL IV outside of the islet in the IM and found a decrease in exocrine COL IV-stained area normalized to total exocrine area in Aab+ (p<0.0001) and T1D donors (p<0.001, Figure 6C) compared to healthy controls. Overall, our results support a role for β-cell-specific expression of MMP-3 and the degradation of both exocrine and endocrine COL IV during T1D pathogenesis in humans.

**Figure 5:**
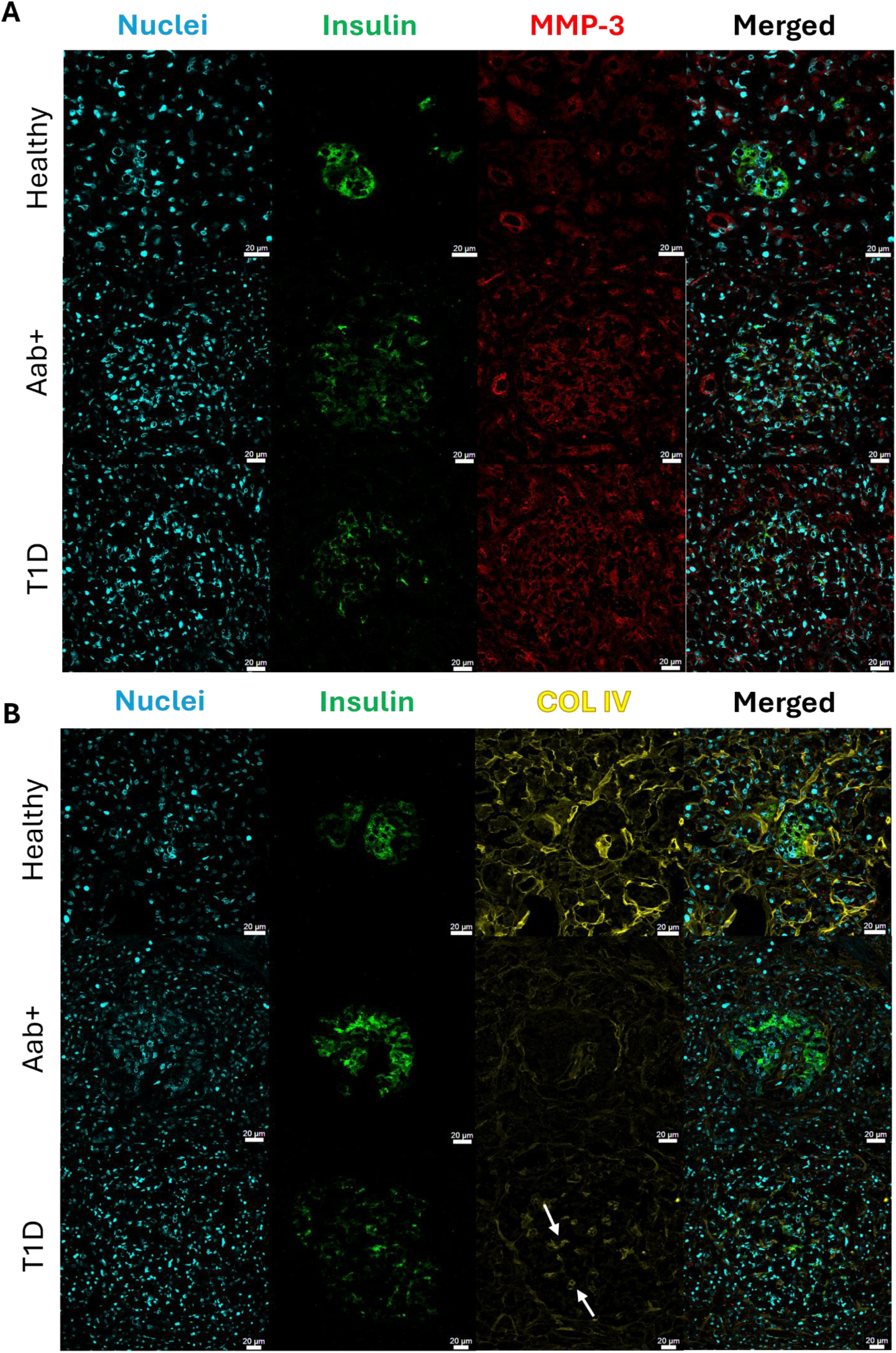
MMP-3 and COL IV IHC analysis in human pancreas sections. Representative confocal microscopy images of (A) MMP-3 and (B) COL IV in healthy, autoantibody positive (Aab+, 2+), and recent onset (<6 months) T1D human donor pancreas sections. MMP-3 and COL IV staining was done on consecutive sections of tissue in each condition. Insulin staining in green was used to identify pancreatic islets. White arrows indicate COL IV staining in islet vasculature. All scale bars are 20μm.

**Figure 6:**
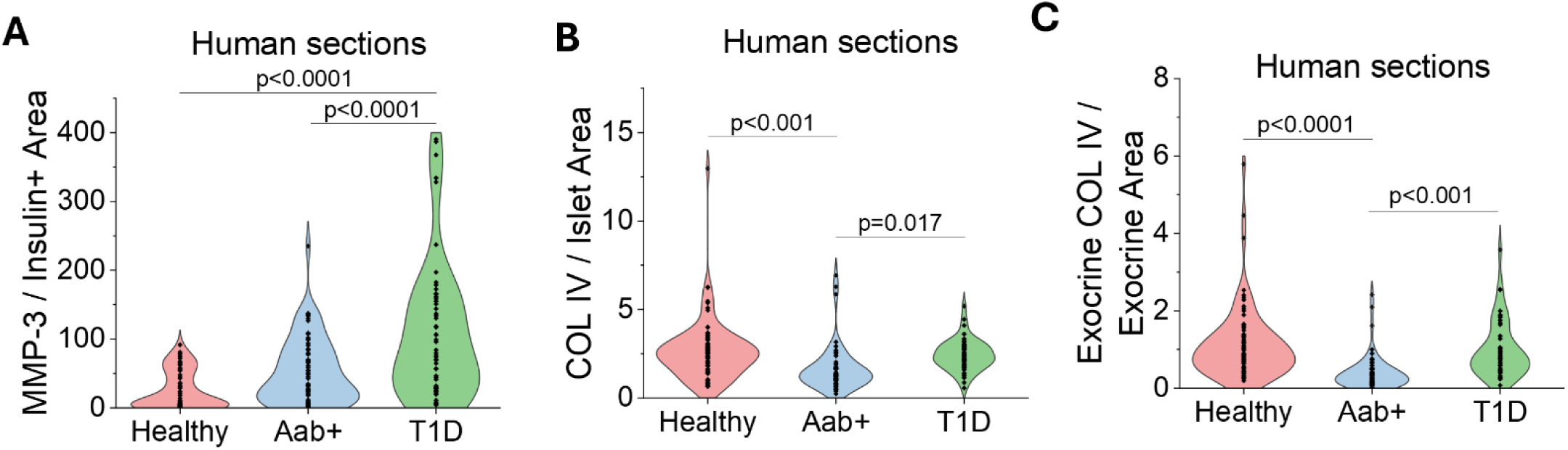
Hyperglycemia and insulitis-induced expression of MMP-3 and degradation of COL IV in human pancreas sections. (A) Quantification of MMP-3 intensity multiplied by staining area normalized to insulin positive area in the confocal images from Figure 5A for healthy, Aab+, and T1D human donor sections. (B) Quantification of COL IV-stained area normalized to total islet area in the confocal images from Figure 5B for healthy, Aab+, and T1D human donor sections. (C) Quantification of exocrine COL IV-stained area normalized to exocrine area in the confocal images from Figure 5B for healthy, Aab+, and T1D human donor sections. p<0.05 indicates statistical significance as determine by ANOVA with Tukey’s post hoc analysis.

### Proinflammatory Cytokine Treatment Influences ECM-related Transcipts in Human Islets

Bulk RNA sequencing data from Wu et al^34^ was previously generated from cadaveric human donor islets treated with or without 50 units/mL IL-1β and 1,000 units/mL IFN-γ for 24h. All raw sequencing counts were obtained from the Gene Expression Omnibus under the accession number GEO GSE169221. Sequencing counts were filtered to remove genes with more than 1 zero read across samples. The fold change in counts of cytokine treated samples compared to untreated controls and p-values were calculated for all donors as well as separate analysis of male and female donors. Additionally, p-values were adjusted using the Benjamini-Hochberg False Discovery Rate (FDR). The log_2_ fold change and the -log_10_ false discovery rate (FDR) adjusted p-value of the data was plotted on a volcano plot for female and male donors (Figure S4). Female donors showed a reduced response to pro-inflammatory cytokines with only 165 differentially expressed transcripts compared to males where 6784 differentially expressed transcripts were identified (Figure S4). In both male and female donors cytokines upregulated transcripts associated with response to inflammation, including GBP1, 2, 4, and 5 (Figure S4). To further determine if cytokine stress upregulates ECM remodeling in human islets, transcripts were filtered using gene ontology (GO) analysis to identify transcripts associated with “extracellular matrix” across all donors (male and female combined) and plotted only these transcripts on a volcano plot (Figure 7A). We identified several significantly upregulated transcripts with function related to collagen catabolism including MMP-1, MMP-3, MMP-10, and MMP-25, with a 3.72, 7.44, 9.52, and 18.75-fold increase in the cytokine treated group compared to the control, respectively (p=0.0088, p=0.0210, p=0.0071, p=0.0005, Figure 7A). Other ECM-modifying proteases include ADAMTS1, ADAMTS4, and ADAMTS9, which all increased 3.41, 4.72, and 3.98-fold, respectively, in the cytokine treated condition compared to the control (p=0.0005, p=0.0035, p=0.0015, Figure 7A). Additionally, transcripts related collagen production were significantly reduced in cytokine treated islets compared to healthy controls, including COL 4A5 (COL IV), COL 26A1, and COL 14A1 with a 0.31, 0.24, and 0.25-fold decrease in expression respectively (p=0.0051, p=0.0028, p=0.0019, Figure 7A). Further analysis of GO results filtered by extracellular matrix related pathways revealed that the top three upregulated biological processes in cytokine treated human islets compared to the control were extracellular matrix organization, cell adhesion, and proteolysis (Figure 7B). Additional upregulated processes include collagen catabolic process, extracellular matrix disassembly, and collagen fibril organization. The following statistically significant transcripts fell into the upregulated extracellular matrix organization and proteolysis biological processes: ADAMTS1, 3, 4, 9, ADAMTS18, MMP-1-3, MMP-9, 10, 12, 19, and MMP-25. The top three downregulated biological processes in cytokine treated human islets compared to the control were extracellular matrix organization, cell adhesion, and collagen fibril organization (Figure 7C). Additional downregulated processes of interest include cell-matrix adhesion and basement membrane organization. The following statistically significant transcripts fell into the downregulated extracellular matrix organization and cell adhesion biological processes: ADAMTS2, 10, 12, 14-17, COL1A1, COL3A1, COL4A1-5, and MMP-7, 11, and MMP-28. Overall, these findings suggest that human islets are capable of remodeling of their ECM in response to stressors such as cytokine exposure, providing strong evidence in support of β-cells degradation of the peri-islet ECM during the progression of pre-clinical T1D.

**Figure 7:**
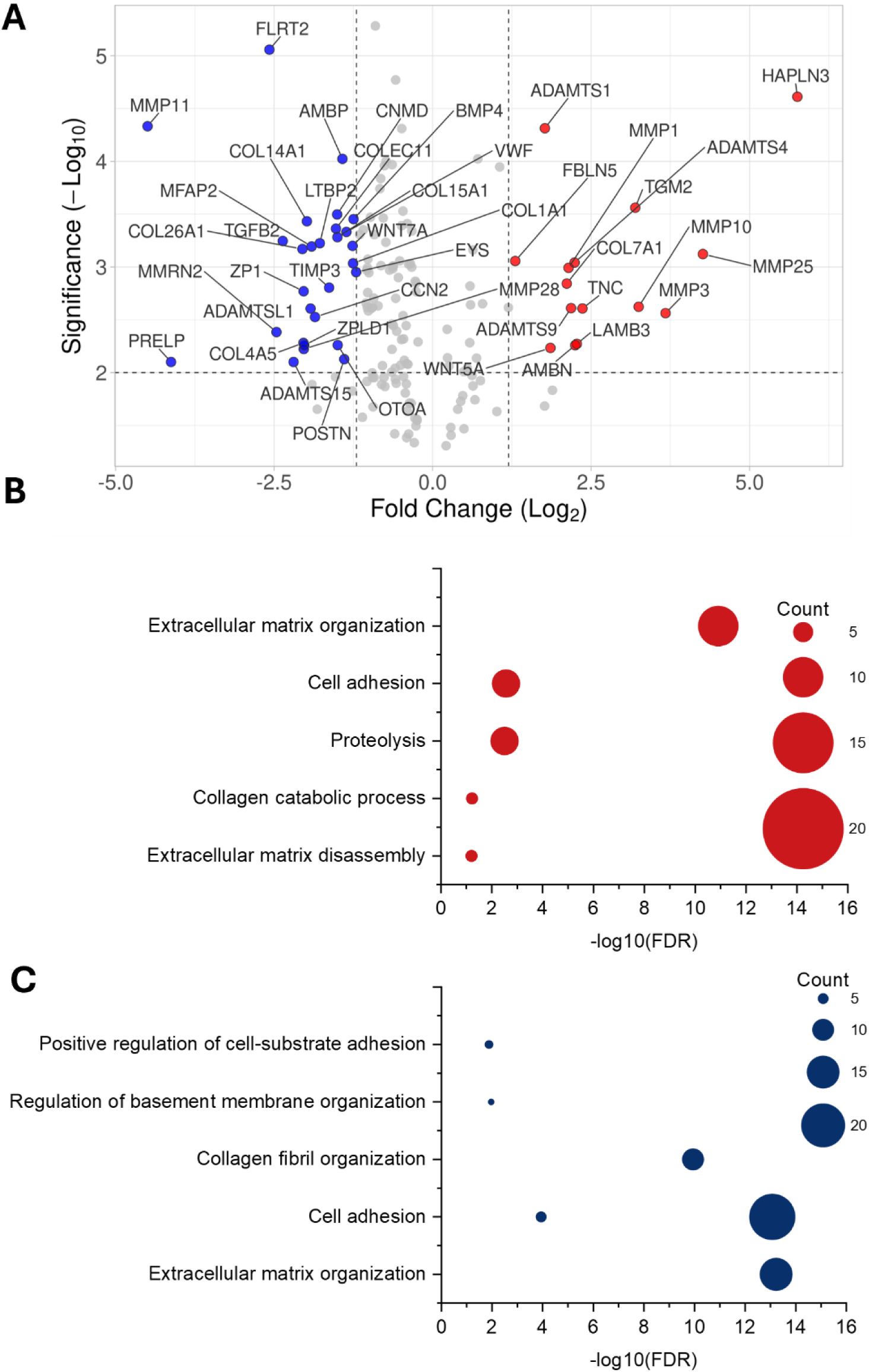
Transcriptomics analysis implicates cytokine treated human islets in ECM degradation. (A) Volcano plot of all statistically significant transcripts associated with changes in the ECM. Transcripts represented by a red dot are upregulated and blue dots are downregulated upon cytokine treatment in human islets. (B) Gene ontology (GO) biological process bubble chart of all statistically significant upregulated proteins in the cytokine treated group relative to the control group. The size of the bubble represents the number of transcripts associated with that GO term. (C) GO biological process bubble chart of all statistically significant downregulated proteins in the cytokine treated group relative to the control group. The GO phrase “extracellular matrix” was used to filter proteins of interest in B and C.

## Discussion

The goal of this study was to determine if stressed β-cells contribute to remodeling and breakdown of the peri-islet ECM during the progression of preclinical T1D. Our study focused primarily on COL IV, one of the primary components of the peri-islet ECM capsule and MMP-3 because of its ability to degrade COL IV and its activation of other MMPs, thus amplifying ECM remodeling in inflammatory conditions^15,18,28,29^. We utilized mouse models of T1D, isolated mouse and human islets exposed to cytokine and hyperglycemic stress, and human donor pancreas from Aab+ or recent onset T1D donors compared to healthy controls. *In vitro*, MMP-3 gene and protein expression increased in both mouse and human islets under cytokine and hyperglycemic stress. In addition, we observed a striking increase in human MMP-3 gene expression during cytokine stress; however, this was not reflected to the same extent at the protein level in cell lysates. We suspect this discrepancy is due to MMP-3 being a secreted matrix metalloproteinase, and thus MMP-3 was likely released into the media in response to cytokine stress or hyperglycemia^35,36^. These results highlight the ability of stressed islets to remodel their environment and contribute to the degradation of the peri-islet BM. This supports a role for the β-cell in maintaining the peri-islet ECM in the healthy pancreas and challenges the current paradigm that infiltrating immune cells are the primary mediator of ECM loss in T1D^4,6^. Additionally, our results support a potential role for stressed β-cells in remodeling the pancreas environment in other pancreatic diseases, such as type 2 diabetes, where significant cytokine and hyperglycemic stress may occur^32^.

In the NOD mouse model of T1D, MMP-3 expression progressively increased from normoglycemia to hyperglycemia, independent of insulitis score. This was accompanied by a corresponding reduction in COL IV, suggesting that COL IV degradation may occur independently of overt immune infiltration because of β-cell MMP-3 secretion. COL IV was preserved in NOD-Scid mice, indicating that autoimmune stress is a key driver of ECM disruption. However, in hyperglycemic NOD mice, COL IV degradation was more extensive and MMP-3 expression was greater than in age-matched normoglycemic counterparts, again independent of insulitis score, implying that hyperglycemia alone may be sufficient to induce ECM degradation by the β-cell. Notably, MMP-3 expression also increased with insulitis severity and was particularly elevated in highly infiltrated islets of hyperglycemic mice, where COL IV loss was most pronounced. Consistent with our findings, previous studies have shown that islet BM degradation coincides with intra-islet immune infiltration, marking the transition from non-destructive to destructive insulitis^3,4,6,37^. These findings suggest that β-cells may upregulate MMP-3 in response to inflammatory cues from infiltrating immune cells, contributing to local ECM breakdown. As the ECM barrier deteriorates, activated immune cells may more readily access insulin-producing β-cells, potentially accelerating β-cell destruction. We believe this provides an explanation for the rapid decline in β-cells from stage 1 to stage 3 of T1D pathogenesis^3,4,12^. The contributions of stressed β-cells to loss of COL IV in the per-islet ECM was particularly evident under conditions of severe immune infiltration and hyperglycemia, suggesting that hyperglycemia may exacerbate ECM degradation when immune activity is already heightened. Collectively, these findings support a multifactorial process of peri-islet COL IV degradation, where inflammatory and hyperglycemic stress on β-cells causes them to compromise ECM integrity, which may contribute to the transition from early to more destructive stages of T1D progression^4^.

Due to the heterogeneous nature of human islet architecture, more rare occurance of insulitis in Aab+ and recent onset T1D donors, and variability in tissue section quality, there was greater variability in insulin and nuclear staining, which presented challenges for our ability to score insulitis analogous to the mouse data. Despite these challenges, MMP-3 and COL IV staining in human pancreas sections revealed trends consistent with those observed in our mouse models where MMP-3 expression increased and COL IV staining decreased in both Aab+ and T1D donors, reinforcing the translational relevance of our findings. Futhermore, our data suggest that β-cells degradation of COL IV occurs in Aab+ donors prior to significant loss of β-cell mass, implicating COL IV degradation as a feature beginning before T1D clinical diagnosis. While several studies have identified a loss of ECM in Aab+ donors, our results provide the first evidence to support a role for the β-cell in contributing to loss of the ECM^3,6,37^. In our human sections, we also observed recovery of COL IV in the islet pervading microvascualture, while the peri-islet COL IV capsule generally remained compromised even in sections with demonstrated recovery of exocrine COL IV. A recent study found that resident pericytes within the islet microvasculature of human donors are active ECM contributors both in physiological and pathophysiological conditions, but in T1D, they contribute enhanced COL IV production in the islet vascualture^38^. This reported pericyte dysfunction supports our observations and may further stress the islet by impairing its ability to regulate blood flow in response to metabolic demands, like during hyperglycemia, potentially compromising insulin secretion. Beyond the islet, exocrine COL IV, including the peri-islet capsule and surrounding IM, also exhibited progressive degradation as disease advanced from healthy to Aab+ and T1D donors. This finding aligns with prior literature documenting a global loss of both peri-islet BM and IM components at sites of immune infiltration and during T1D in humans^3,6^. Islet interactions with the ECM provide crititcal cues to support islet survival that when lost during T1D may make islets more suseptible to immune-mediated killing^10^. Together, these observations highlight that stressed β-cells remodel the peri-islet ECM and that expression of MMP-3 throughout the development of T1D contributes to loss of COL IV in the peri-islet ECM that may accelerate immune infiltration and β-cell death.

Transcriptomics analysis of cytokine-treated human islets revealed the upregulation of several ECM-degrading enzymes, including MMPs and ADAMTS family members, which target a broad range of BM and IM components^30,39^. Together with our previous findings, these results suggest that inflammatory signals may trigger a transcriptional program within islets that promotes ECM remodeling. Significant sex-based differences were observed in response to cytokines in male versus female donors, where female donors were more resistant to inflammation induced transcriptional changes compared to males. This is consistent with previous studies that found islets from female donors are more resistant to dysfunction and ER stress in type 2 diabetes^40^. While our transcriptomics analysis was underpowered (n=4 compared to n=6 for males) to detect significantly altered transcripts related to ECM remodeling in female donors, our predominantly female nPOD Aab+ donors support similar ECM degradation and β-cell specific expression of MMP-3 in males and females. While many of the upregulated transcripts (sex exclusive) are associated with collagen remodeling, several transcripts, including members of the ADAMTS family that target proteoglycans (aggrecan and versican), glycoproteins, and laminin in addition to collagen, provide evidence for broad β-cell remodeling of the peri-islet microenvironment^39^. Additionally, MMP-10 and MMP-12 which target fibronectin and laminin respectively, were found to be upregualted and have been shown to activate other MMPs that may promote further remodeling of the ECM^18^. Overall, these changes in ECM remodeling transcripts support a scenario in which stressed β-cells compromise the structural integrity of the ECM in early stages of T1D, potentially compounding immune infiltration into the islet. Conversely, downregulated transcripts included collagens critical to BM and IM structure, as well as ECM remodeling enzymes that are less degradative or are primarily involved in matrix assembly and maintenance. For instance, ADAMTS2 and ADAMTS14 are known to contribute to collagen maturation and fibril stabilization through their roles in procollagen processing^39,41^. MMP-7, MMP-11, and MMP-28 are associated with tissue repair and wound healing, and are not typically categorized as major ECM-degrading enzymes^18,30^. Their downregulation suggests a reduction in ECM repair or synthesis capacity, potentially limiting the islet’s ability to preserve or restore its protective niche. However, this pro-degradative phenotype may shift in recent onset or longstanding T1D where expression of these MMPs may shift β-cells to a pro-repair phenotype that could contribute to the increase in exocrine COL IV that was observed in T1D donors compared to Aab+ donors. Additional studies with T1D donors are needed to confirm this hypothesis. Altogether, these data suggest that cytokine stress drives destructive ECM remodeling by islets, weakening the local microenvironment and potentially accelerating β-cell loss during T1D pathogenesis. Future studies should aim to further define the expression of specific ECM-degrading enzymes from stressed islets and to determine how their expression and activity are regulated in T1D.

In summary, the results of our study are supported by previous studies that have demonstrated global loss of key ECM components at sites of immune infiltration in human T1D donor samples^3,4,6,20^. Our study not only reinforces these observations but also builds upon prior work to define several newly identified events in the pathogenesis of T1D. First, we provide direct evidence that stressed β-cells themselves contribute to ECM remodeling by upregulation of MMP-3 and a host of other ECM degrading enzymes, suggesting that β-cells aid in their own demise. This supports and expands upon prior hypotheses that β-cells may actively participate in creating a permissive environment for immune invasion^24^. Second, our findings demonstrate that ECM degradation begins during the preclinical stages of T1D and that β-cell contributions to ECM degradation are independent of immune infiltration degree. Additionally, we demonstrate that recovery of COL IV is limited to the vasculature and to a lesser extent the exocrine pancreas and is generally not recovered in the peri-islet ECM capsule in patients with recent onset T1D. Overall, our data demonstrates that stressed β-cells are contributing to the remodeling of the peri-islet ECM, specifically the degradation of COL IV. We postulate that the degradation of this protective barrier reduces pro-survival signaling from the ECM to the islet and permits a rapid influx of activated immune cells, thereby accelerating the decline of insulin-producing β-cells during T1D pathogenesis. By clarifying the spatial and temporal dynamics of ECM degradation, as well as the role of stressed β-cells in this process, we gain a deeper understanding of peri-islet ECM barrier loss during T1D pathogenesis. This insight, along with future studies, could be leveraged to develop therapeutic strategies aimed at preserving ECM integrity and preventing immune infiltration, ultimately protecting β-cells before the rapid decline that leads to clinical diagnosis of T1D.

## Materials & Methods

All NOD mice used were female due to their higher incidence rate of T1D compared to males. Additionally, the limited availability of T1D donor pancreas and high value of human cadaveric islets and tissues precluded stratification based on donor sex. Transcriptomics data was stratified based on donor sex to determine if there were sex-based differences in response to cytokine stress.

### 1. Animal care

Mice were housed in a temperature- and light-controlled environment with 12-hour light-dark cycles, and they were provided with access to food and drink ad libitum. C57Bl/6, NOD, and NOD-Scid mice were purchased from the Jackson Laboratories (strain #000664) at 8 weeks of age.

### 2. Human islets

Human islets were obtained from the Integrated Islet Distribution Program (IIDP) from the following donors as outlined in Table 1:

**Table 1:** Human cadaveric islet donor demographics and isolated islet viability and purity for islets obtained through the Integrated Islet Distribution Program (IIDP).

| RRID# | Viability (%) | Purity (%) | Age | Sex | Race/Ethnicity |
| --- | --- | --- | --- | --- | --- |
| SAMN43474066 | 95 | 95 | 54 | Male | Not documented/Hispanic or Latino |
| SAMN43463688 | 95 | 90 | 54 | Male | Black or African American/Not Hispanic or Latino |
| SAMN43897604 | 95 | 90 | 41 | Male | Other/Hispanic or Latino |
| SAMN45149991 | 95 | 94 | 66 | Male | White/Not Hispanic or Latino |
| SAMN47416577 | 95 | 90 | 56 | Male | White/Not Hispanic or Latino |

Human donor tissue is de-identified through the IIDP and is therefore exempt from requiring human subjects research protocols. Islets that were clear of any excess pancreatic tissue and were consistent in color and shape based on inspection through a microscope were selected for use in experiments.

### 3. Islet isolation and culture

Islets were isolated from 8–16-week-old female C57BL/6 mice, 12-week-old female normoglycemic NOD mice, and 12-week-old female NOD-Scid mice. NOD mouse diabetes progression was monitored by weekly ad lib glucose measurements taken from the tail vain. Mice with a blood glucose level >250 mg/dl for three consecutive measurements were considered hyperglycemic and euthanized. Animals were injected with 100mg/kg ketamine and 8mg/kg xylazine and euthanized via exsanguination. Islets were isolated by injecting the pancreas with 12.5mg/mL collagenase, pancreas removal, and enzymatic digestion at 37°C. Islets were handpicked into 1640 RPMI Medium with 1X with L-glutamine and 25 mM HEPES (Fisher Scientific, Hampton, NH) with 10% fetal bovine serum (Fisher Scientific, Hampton, NH), 10,000 U/mL Penicillin and 10,000μg/mL Streptomycin (Sigma-Aldrich, St. Louis, MO) and incubated at 37°C and 5% CO_2_ for a minimum of 3 hours before commencing experiments. Islets that were clear of any excess pancreatic tissue and were consistent in color and shape based on inspection through a microscope were selected for use in experiments.

### 4. Quantitative polymerase chain reaction (qPCR)

C57BL/6 mouse islets and human islets were treated with or without a 0.1X (1ng/ml TNF-α, 0.5ng/ml IL-1β, 10ng/ml IFN-γ) proinflammatory cytokine cocktail for 48h and 72h. This cytokine cocktail was chosen for its synergistic effects on islet dysfunction and death, as well as its high abundance in the T1D pancreatic environment^26,31,42^. All qPCR samples were purified for mRNA by using RNeasy® Mini kit (74104; Qiagen, Germantown, MA) following the manufacturers’ instruction. Reverse transcription was performed on mRNA samples using Applied Biosystems™ High-Capacity cDNA Reverse Transcription Kit (43-688-14; Fisher Scientific, Hampton, NH) following the manufacturers’ instruction. The resulting cDNA was used to analyze the expression of MMP-3 and housekeeping gene HPRT1 by real-time quantitative PCR (Roche 480) in 10-μL reactions containing 2X Applied Biosystems™ PowerUp™ SYBR™ Green Master Mix (A25776; Fisher Scientific, Hampton, NH) and 20μM primers. Table 2 lists all primer sequences used:

**Table 2:**
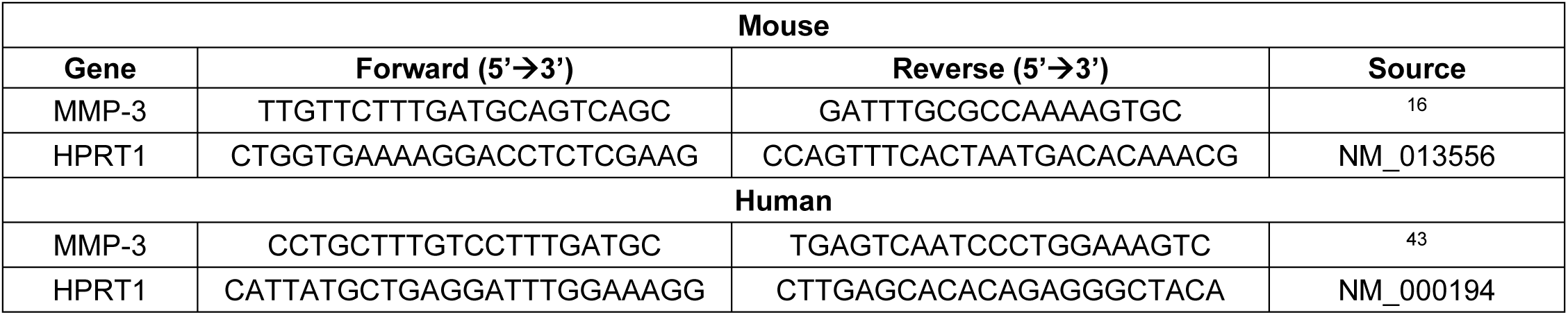
qPCR primer sequences for mouse and human experiments.

All qPCR primers were purchased from Integrated DNA Technologies (Coralville, IA). The results were calculated using Eqn. 1-3 below (n=3).

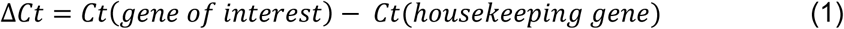

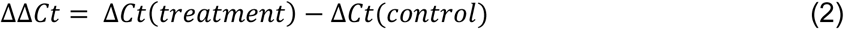

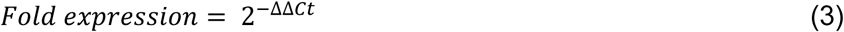

### 5. Western blotting

C57BL/6 mouse islets and human islets were treated with or without a 0.1X (1ng/ml TNF-α, 0.5ng/ml IL-1β, 10ng/ml IFN-γ) or 1X (10ng/ml TNF-α, 5ng/ml IL-1β, 100ng/ml IFN-γ) proinflammatory cytokine cocktail for 24, 48, and 72h. Islets were collected and lysed in lysis buffer containing 1X protease and phosphatase inhibitors (ThermoFisher Scientific, Waltham, MA). Protein content was measured using the Pierce BCA Protein Assay Kit (PI23225; Fisher Scientific, Hampton, NH) according to the manufacturer’s instructions. Samples were run on 4–15 % mini-PROTEAN® TGX protein gels (Bio-Rad, Hercules, CA) and transferred to a PVDF (Azure Biosystems, Dublin, CA) membrane. PVDF membranes were blocked in chemi-blot blocking buffer (Azure Biosystems, Dublin, CA) for 2hr and probed with anti-MMP-3 (17873-1-AP; Proteintech, Rosemont, IL) at a dilution of 1:500 for >12hr at 4°C. Blots were washed in washing buffer (1X PBS with 0.1 % Tween) 3x prior to the addition of secondary anti-rabbit (102649–670; VWR, Radnor, PA) horseradish peroxidase-conjugated antibody diluted to 1:10,000 for 2hr at room temperature. The membranes were washed in washing buffer 3x and incubated with Radiance Plus (Azure Biosystems, Dublin, CA) for 2 min in the dark. All samples were normalized to protein content by probing with anti-β-actin (sc-47778; Santa Cruz Biotechnology, Santa Cruz, CA) at a dilution of 1:100 for >12hr at 4°C. Blots were washed in washing buffer (1X PBS with 0.1 % Tween) 3x prior to the addition of secondary anti-mouse (626520; Fisher Scientific, Hampton, NH) horseradish peroxidase-conjugated antibody diluted to 1:1000 for 2hr at room temperature. All membranes were imaged using an Azure c600 imaging system (Azure Biosystems, Dublin, CA) and protein quantification was performed in ImageJ using densitometric analysis (n=3).

### 6. Immunohistochemistry (IHC)

Frozen, mouse tissue blocks were sectioned at 10-µm intervals. For each pancreas, a total of 45 consecutive serial sections were obtained. Three consecutive slices of mouse tissue were placed on each slide. Frozen sections of human pancreas were provided by the network of Pancreatic Organ Donors (nPOD). Characteristics of the donors examined are presented in Table 3:

**Table 3:** Clinical characteristics of controls, AA+ patients, and patients with type 1 diabetes.

| Donor ID | Donor Type | Aab+ | Age | Gender | Ethnicity | HbA1c | BMI |
| --- | --- | --- | --- | --- | --- | --- | --- |
| 6521 | Aab+ | GADA / IA2A / ZnT8A | 19.77 | M | Hispanic | 5.8 | 24.1 |
| 6450 | Aab+ | GADA / ZnT8 | 22 | F | Caucasian | 5.7 | 24.4 |
| 6267 | Aab+ | GADA / IA2A | 23 | F | Caucasian | 5 | 23.5 |
| 6414 | T1D | GADA / mIAA / ZnT8 | 23.1 | M | Caucasian | 14 | 28.4 |
| 6362 | T1D | GADA | 24.9 | M | Caucasian | 10 | 28.5 |
| 6578 | T1D | IA2A / ZnT8 | 11.95 | F | Caucasian | 13.6 | 22.5 |
| 6546 | No diabetes |  | 22.29 | M | Asian |  | 23.7 |
| 6605 | No diabetes |  | 23.2 | F | Hispanic |  | 22.7 |
| 6227 | No diabetes |  | 17 | F | Caucasian |  | 26.4 |

Tissue sections were defrosted, and a hydrophobic border was drawn around each tissue slice using a barrier pen. Sections were permeabilized with permeabilization buffer (5% normal donkey serum (NDS) and 0.25% Triton-X-100 in 1X phosphate-buffered saline (PBS)) for 5 min and blocked with blocking buffer (5% NDS and 0.1% Triton-X-100 in 1X PBS) for 5 min at room temperature. Sections were incubated overnight at 4°C with a primary antibody combination of (1) anti-mouse insulin-Alexa 488 (1:100, 50-112-4642; Fisher Scientific, Hampton, NH) and anti-rabbit MMP-3 (1:100, 17873-1-AP; Proteintech, Rosemont, IL) or (2) anti-mouse insulin-Alexa 488 (1:100, 50-112-4642; Fisher Scientific, Hampton, NH), anti-rabbit collagen IV (1:500, ab6586; Abcam, Waltham, MA), and anti-rat CD3+ (1:100, 14-0032-82; Fisher Scientific, Hampton, NH). After incubation, the slides were rinsed with 1X PBS and incubated with appropriate secondary antibodies (goat anti-rabbit Alexa 568 (1:100, A11011; ThermoFisher Scientific, Waltham, MA) for MMP-3 and collagen IV and goat anti-rat Alexa 647 (1:100, A21247; ThermoFisher Scientific, Waltham, MA) for CD3+) for 2h covered with foil at room temperature. All the slides were rinsed with 1X PBS, mounted with fluoromount containing DAPI, and sealed for imaging.

Sections were imaged on a Leica STELLARIS 5 Confocal Microscope with LIAchroic laser supply unit with a 40X water immersion objective using 405 nm, 488 nm, 514 nm, 647 nm solid state lasers and HyD spectral detectors. To analyze the amount of MMP-3 and COL IV expression in pancreas sections from NOD/NOD-Scid mice and human donor pancreas, IHC images were analyzed semi-manually in Fiji (ImageJ, NIH). A representative image of MMP 3 staining is shown in Figure S3A, where the insulin positive area was outlined and the background was cleared (Figure S3B). Next, the channels were split and the MMP-3 channel was selected (Figure S3C). The 3D Objects Counter in ImageJ was used to identify MMP-3 positive areas using a consistent threshold for determining positive signal for all channels. An example of the positive areas after threshold was selected can be seen in Figure S3D. The total surface area, mean fluorescence intensity, and background fluorescence intensity were recorded. Analysis of COL IV positive areas was the same as for MMP-3 positive areas except for outlining the islet area which was determined by the presence of the peri-islet COL IV capsule rather than only insulin positive areas (Figure S3E-H). The same process for insulin was repeated to obtain insulin positive areas. MMP-3 and COL IV signal was calculated by multiplying the total MMP-3 positive area by the average fluorescence intensity and dividing by the total insulin positive area.

The insulitis score for each islet was recorded for NOD samples based on CD3+ staining and density of nuclei within the islet; however, insulitis scoring was not possible for the human samples as infiltration with CD3+ cells is a rare event in human islets despite confirmation of insulitis in the tissue blocks utilized for this study. For NOD islets, an islet was assigned a score of 0 if no infiltration was observed, a score of 1 if only a thin ring of infiltration outlined the islet, a score of 2 if less than 50% of the islet was infiltrated, and a score of 3 if more than 50% of the islet was infiltrated.

### 7. Transcriptomics

RNA sequencing data was obtained from Wu et al^34^. Briefly, human islets were obtained from the Integrated Islet Distribution Program (IIDP) or the University of Alberta from the following donors as outlined in Table 4. Islets were cultured overnight in standard Prodo medium (Prodo Laboratories, Aliso Viejo, CA) and treated with or without 50 units/mL IL-1β and 1,000 units/mL IFN-γ for 24h.

**Table 4:** Human cadaveric islet donor demographics and isolated islet viability and purity for islets obtained through the Integrated Islet Distribution Program (IIDP) or the University of Alberta for RNA sequencing.

| ID | Gender | Age | BMI | Islet purity (%) | Islet viability (%) |
| --- | --- | --- | --- | --- | --- |
| 1 | Female | 33 | 34.2 | 95 | 98 |
| 2 | Female | 43 | 29.5 | 80 | 96 |
| 3 | Male | 35 | 24.2 | 95 | 95 |
| 4 | Male | 30 | 35.9 | 90 | 95 |
| 5 | Female | 52 | 25.9 | 90 | 95 |
| 6 | Male | 23 | 24.8 | 95 | 95 |
| 7 | Female | 44 | 34.5 | 90 | 90 |
| 8 | Male | 28 | 24.6 | 91 | 95 |
| 9 | Male | 15 | 24.6 | 80 | 97 |
| 10 | Male | 28 | 30.7 | 95 | 95 |

Raw sequence data was obtained from the Gene Expression Omnibus under the accession number GEO GSE169221. The log_2_ fold change and the -log_10_ false discovery rate (FDR) adjusted p-value of the raw data was calculated and plotted into a volcano plot using VolcaNoseR (https://huygens.science.uva.nl/VolcaNoseR2/). Statistical significance was defined as an FDR p-value ≤ 0.05 for comparison between cytokine-treated and untreated, control human islets. Gene Ontology (GO) enrichment analysis was performed with DAVID (https://david.ncifcrf.gov/tools.jsp) using the biological processes category. The GO phrase “extracellular matrix” was used to further filter proteins of interest.

### 8. Statistical Analysis

Data represents the average over all islets for each measurement with error bars representing standard error unless otherwise noted. Statistics were performed using Origin software (OriginLabs, Northampton, MA). Two sample t-test, one-way and two-way ANOVA with Tukey’s post hoc analysis were performed as indicated. A p-value of <0.05 or 95% confidence intervals were considered statistically significant unless otherwise noted.

### 9. Study Approval

All experiments using mice were performed at the University of Colorado Anschutz Medical Campus and in compliance with the guidelines and relevant laws set by the University of Colorado and the National Institutes of Health guide for the care and use of Laboratory animals. All performed procedures were approved by the University of Colorado Institutional Animal Care and Use Committee (Protocol 00929).

### 10. Data Availability

All images of stained human pancreas samples will be available by request through the nPOD data repository. Data from qPCR, western blot, and IHC quantification will be available by request to the corresponding author. Image analysis scripts are freely available for download and use with ImageJ (https://imagej.net/ij/).

## Author contributions

N.L.F. conceptualized and received funding for the project and oversaw experiments and data analysis. N.L.F. and C.G.J wrote and edited the manuscript. C.G.J. and K.L. performed experiments. C.G.J., K.L., and N.L.F. analyzed data. All authors have given approval to the final version of the manuscript.

## Acknowledgments

The authors would like to acknowledge the funding sources that made this work possible, including the following grants: American Diabetes Association grant 7-21-JDF-020 to NLF, NIDDK grant F31 DK132926 to C.M.J., NIDDK grant 5R01DK137221 to NLF, and Pilot Funding through the Helmsley Charitable Trust George S. Eisenbarth nPOD Award for Team Science to NLF. Additionally, the Farnsworth lab is supported by funding from Breakthrough T1D (formerly JDRF) 3-SRA-2023-1367-S-B. This research was performed with the support of the

Network for Pancreatic Organ donors with Diabetes (nPOD; RRID:SCR_014641), a collaborative type 1 diabetes research project supported by Breakthrough T1D and The Leona M. & Harry B. Helmsley Charitable Trust (Grant#3-SRA-2023-1417-S-B). The content and views expressed are the responsibility of the authors and do not necessarily reflect the official view of nPOD or the National Institutes of Health. Organ Procurement Organizations (OPO) partnering with nPOD to provide research resources are listed at https://npod.org/for-partners/npod-partners/. Experiments with isolated islets utilized the Islet Isolation Core in the University of Colorado Diabetes Research Center funded by NIDDK grant # P30-DK116073. Additionally, we would like to thank the organ donors and their families for their generous gift that has made this work possible.

## References

(1) Stendahl, J. C.; Kaufman, D. B.; Stupp, S. I. Extracellular Matrix in Pancreatic Islets: Relevance to Scaffold Design and Transplantation. Cell Transplant. 2009, 18 (1), 1–12. 10.3727/096368909788237195.

(2) Alessandra, G.; Algerta, M.; Paola, M.; Carsten, S.; Cristina, L.; Paolo, M.; Gabriella, T.; Carla, P. Shaping Pancreatic B-Cell Differentiation and Functioning : The Influence of Mechanotransduction. Cells 2020, 9 (413), 1–24.

(3) Korpos, É.; Kadri, N.; Kappelhoff, R.; Wegner, J.; Overall, C. M.; Weber, E.; Holmberg, D.; Cardell, S.; Sorokin, L. The Peri-Islet Basement Membrane, a Barrier to Infiltrating Leukocytes in Type 1 Diabetes in Mouse and Human. Diabetes 2013, 62 (2), 531–542. 10.2337/db12-0432.

(4) Irving-Rodgers, H. F.; Ziolkowski, A. F.; Parish, C. R.; Sado, Y.; Ninomiya, Y.; Simeonovic, C. J.; Rodgers, R. J. Molecular Composition of the Peri-Islet Basement Membrane in NOD Mice: A Barrier against Destructive Insulitis. Diabetologia 2008, 51 (9), 1680–1688. 10.1007/s00125-008-1085-x.

(5) Ziolkowski, A. F.; Popp, S. K.; Freeman, C.; Parish, C. R.; Simeonovic, C. J. Heparan Sulfate and Heparanase Play Key Roles in Mouse β Cell Survival and Autoimmune Diabetes. J. Clin. Invest. 2012, 122 (1), 132–141. 10.1172/JCI46177.

(6) Bogdani, M.; Korpos, E.; Simeonovic, Charmaine, J.; Parish, Christopher, R.; Sorokin, L.; Wight, Thomas, N. Extracellular Matrix Components in the Pathogenesis of Type 1 Diabetes. Curr Diab Rep 2014, 14 (552), 1–11. 10.1161/CIRCULATIONAHA.110.956839.

(7) Tanzer, M. L. Current Concepts of Extracellular Matrix. J. Orthop. Sci. 2006, 11 (3), 326–331. 10.1007/s00776-006-1012-2.

(8) Hynes, R. O. Extracellular Matrix: Not Just Pretty Fibrils. Science (80-. ). 2009, 326 (5957), 1216–1219. 10.1126/science.1176009.Extracellular.

(9) Weber, L. M.; Anseth, K. S. Hydrogel Encapsulation Environments Functionalized with Extracellular Matrix Interactions Increase Islet Insulin Secretion. Matrix Biol. 2008, 27 (8), 667–673. 10.1016/j.matbio.2008.08.001.

(10) Hammar, E.; Parnaud, G.; Bosco, D.; Perriraz, N.; Maedler, K.; Donath, M.; Rouiller, D. G.; Halban, P. A. Extracellular Matrix Protects Pancreatic β-Cells against Apoptosis: Role of Short- and Long-Term Signaling Pathways. Diabetes 2004, 53 (8), 2034–2041. 10.2337/diabetes.53.8.2034.

(11) Child, A.; Larkin, E. J.; Fontaine, M. J. Co-Encapsulation of ECM Proteins to Enhance Pancreatic Islet Cell Function; Elsevier Inc., 2019; Vol. 2. 10.1016/B978-0-12-814831-0.00022-1.

(12) Powers, A. C. Type 1 Diabetes Mellitus: Much Progress, Many Opportunities. J. Clin. Invest. 2021, 131(8), 1–10. 10.1172/JCI142242.

(13) Katsarou, A.; Gudbjörnsdottir, S.; Rawshani, A.; Dabelea, D.; Bonifacio, E.; Anderson, B. J.; Jacobsen, L. M.; Schatz, D. A.; Lernmark, A. Type 1 Diabetes Mellitus. Nat. Rev. Dis. Prim. 2017, 3, 1–18. 10.1038/nrdp.2017.16.

(14) Achenbach, P.; Bonifacio, E.; Koczwara, K.; Ziegler, A. G. Natural History of Type 1 Diabetes. Diabetes 2005, 54 (SUPPL. 2). 10.2337/diabetes.54.suppl_2.S25.

(15) Jabłońska-Trypuć, A.; Matejczyk, M.; Rosochacki, S. Matrix Metalloproteinases (MMPs), the Main Extracellular Matrix (ECM) Enzymes in Collagen Degradation, as a Target for Anticancer Drugs. J. Enzyme Inhib. Med. Chem. 2016, 31, 177–183. 10.3109/14756366.2016.1161620.

(16) Jehan, F.; Zarka, M.; de la Houssaye, G.; Veziers, J.; Ostertag, A.; Cohen-Solal, M.; Geoffroy, V. New Insights into the Role of Matrix Metalloproteinase 3 (MMP3) in Bone. FASEB BioAdvances 2022, 4 (8), 524–538. 10.1096/fba.2021-00092.

(17) Alper, M.; Kockar, F. IL-6 Upregulates a Disintegrin and Metalloproteinase with Thrombospondin Motifs 2 (ADAMTS-2) in Human Osteosarcoma Cells Mediated by JNK Pathway. Mol. Cell. Biochem. 2014, 393 (1–2), 165–175. 10.1007/s11010-014-2056-9.

(18) Laronha, H.; Caldeira, J. Structure and Function of Human Matrix Metalloproteinases. Cells 2020, 1–18.

(19) Medina, C. O.; Nagy, N.; Bollyky, P. L. Extracellular Matrix and the Maintenance and Loss of Peripheral Immune Tolerance in Autoimmune Insulitis. Curr. Opin. Immunol. 2018, 55, 22–30. 10.1016/j.coi.2018.09.006.

(20) Vaday, G. G.; Lider, O. Extracellular Matrix Moieties, Cytokines, and Enzymes: Dynamic Effects on Immune Cell Behavior and Inflammation. J. Leukoc. Biol. 2000, 67 (2), 149–159. 10.1002/jlb.67.2.149.

(21) Montgomery, A. M. P.; Sabzevari, H.; Reisfeld, R. A. Production and Regulation of Gelatinase B by Human T-Cells. BBA - Mol. Cell Res. 1993, 1176 (3), 265–268. 10.1016/0167-4889(93)90054-S.

(22) Owen, C. A.; Campbell, E. J. The Cell Biology of Leukocyte-Mediated Proteolysis. J. Leukoc. Biol. 1999, 65 (2), 137–150. 10.1002/jlb.65.2.137.

(23) Zhou, H.; Bernhard, E. J.; Fox, F. E.; Billings, P. C. Induction of Metalloproteinase Activity in Human T-Lymphocytes. BBA - Mol. Cell Res. 1993, 1177 (2), 174–178. 10.1016/0167-4889(93)90037-P.

(24) Saunders, D. C.; Aamodt, K. I.; Richardson, T. M.; Hopkirk, A.; Aramandla, R.; Poffenberger, G.; Jenkins, R.; Flaherty, D. K.; Prasad, N.; Levy, S. E.; Powers, A. C.; Brissova, M. Coordinated Interactions between Endothelial Cells and Macrophages in the Islet Microenvironment Promote β Cell Regeneration. bioRxiv 2020, 2020.08.28.265728. 10.1038/s41536-021-00129-z.

(25) Parish, C. R.; Freeman, C.; Ziolkowski, A. F.; He, Y. Q.; Sutcliffe, E. L.; Zafar, A.; Rao, S.; Simeonovic, C. J. Unexpected New Roles for Heparanase in Type 1 Diabetes and Immune Gene Regulation. Matrix Biol. 2013, 32 (5), 228–233. 10.1016/j.matbio.2013.02.007.

(26) Lu, J.; Liu, J.; Li, L.; Lan, Y.; Liang, Y. Cytokines in Type 1 Diabetes: Mechanisms of Action and Immunotherapeutic Targets. Clin. Transl. Immunol. 2020, 9 (3), 1–17. 10.1002/cti2.1122.

(27) Padgett, L. E.; Broniowska, K. A.; Hansen, P. A.; Corbett, J. A.; Tse, H. M. The Role of Reactive Oxygen Species and Proinflammatory Cytokines in Type 1 Diabetes Pathogenesis. Ann. N. Y. Acad. Sci. 2013, 1281, 16–35. 10.1111/j.1749-6632.2012.06826.x.

(28) Lu, P.; Takai, K.; Weaver, V. M.; Werb, Z. Extracellular Matrix Degradation and Remodeling in Development and Disease. Cold Spring Harb. Perspect. Biol. 2011, 3 (12), 1–24. 10.1101/cshperspect.a005058.

(29) Wan, J.; Zhang, G.; Li, X.; Qiu, X.; Ouyang, J.; Dai, J.; Min, S. Matrix Metalloproteinase 3: A Promoting and Destabilizing Factor in the Pathogenesis of Disease and Cell Differentiation. Front. Physiol. 2021, 12 (July), 1–10. 10.3389/fphys.2021.663978.

(30) Chang, M. Matrix Metalloproteinase Profiling and Their Roles in Disease. RSC Adv. 2023, 13 (9), 6304– 6316. 10.1039/d2ra07005g.

(31) Farnsworth, N. L.; Walter, R. L.; Hemmati, A.; Westacott, M. J.; Benninger, R. K. P. Low Level Pro-Inflammatory Cytokines Decrease Connexin36 Gap Junction Coupling in Mouse and Human Islets through Nitric Oxide-Mediated Protein Kinase Cδ. J. Biol. Chem. 2016, 291 (7), 3184–3196. 10.1074/jbc.M115.679506.

(32) Eizirik, D. L.; Korbutt, G. S.; Hellerström, C. Prolonged Exposure of Human Pancreatic Islets to High Glucose Concentrations in Vitro Impairs the β-Cell Function. J. Clin. Invest. 1992, 90 (4), 1263–1268. 10.1172/JCI115989.

(33) Bertuzzi, F.; Saccomanno, K.; Socci, C.; Davalli, A. M.; Taglietti, M. V.; Berra, C.; Dalcin, E.; Monti, L. D.; Pozza, G.; Pontiroli, A. E. Long-Term in Vitro Exposure to High Glucose Increases Proinsulin-like-Molecules Release by Isolated Human Islets. J. Endocrinol. 1998, 158 (2), 205–211. 10.1677/joe.0.1580205.

(34) Wu, W.; Syed, F.; Simpson, E.; Lee, C. C.; Liu, J.; Chang, G.; Dong, C.; Seitz, C.; Eizirik, D. L.; Mirmira, R. G.; Liu, Y.; Evans-Molina, C. Impact of Proinflammatory Cytokines on Alternative Splicing Patterns in Human Islets. Diabetes 2022, 71 (1), 116–127. 10.2337/db20-0847.

(35) Shao, L.; Liu, W.; Zhang, C.; Ma, W.; Yu, X.; Han, J.; Wang, X. The Role and Function of Secretory Protein Matrix Metalloproteinase-3 (MMP3) in Cervical Cancer. Iran. J. Public Health 2024, 53 (4), 855– 866. 10.18502/ijph.v53i4.15562.

(36) Tandara, A. A.; Mustoe, T. A. MMP- and TIMP-Secretion by Human Cutaneous Keratinocytes and Fibroblasts - Impact of Coculture and Hydration. J. Plast. Reconstr. Aesthetic Surg. 2011, 64 (1), 108–116. 10.1016/j.bjps.2010.03.051.

(37) Pavin, E. J.; Pinto, G. A.; Zollner, R. L.; Vassallo, J. Immunohistochemical Study of the Pancreatic Basement Membrane in Non Obese Diabetic Mice (NOD) with Spontaneous Autoimmune Insulitis. J Submicrosc Cytol Pathol. 2003, 35 (1), 25–27.

(38) Mateus Gonçalves, L.; Fahd Qadir, M. M.; Boulina, M.; Makhmutova, M.; Pereira, E.; Almaça, J. Pericyte Dysfunction and Impaired Vasomotion Are Hallmarks of Islets during the Pathogenesis of Type 1 Diabetes. Cell Rep. 2023, 42 (8). 10.1016/j.celrep.2023.112913.

(39) Tang, B. L. ADAMTS: A Novel Family of Extracellular Matrix Proteases. Int. J. Biochem. Cell Biol. 2001, 33 (1), 33–44. 10.1016/S1357-2725(00)00061-3.

(40) Brownrigg, G. P.; Xia, Y. H.; Chu, C. M. J.; Wang, S.; Chao, C.; Zhang, J. A.; Skovsø, S.; Panzhinskiy, E.; Hu, X.; Johnson, J. D.; Rideout, E. J. Sex Differences in Islet Stress Responses Support Female β Cell Resilience. Mol. Metab. 2023, 69 (January), 101678. 10.1016/j.molmet.2023.101678.

(41) Satz-Jacobowitz, B.; Hubmacher, D. The Quest for Substrates and Binding Partners: A Critical Barrier for Understanding the Role of ADAMTS Proteases in Musculoskeletal Development and Disease. Dev. Dyn. 2021, 250 (1), 8–26. 10.1002/dvdy.248.

(42) Rabinovitch, A.; Suarez-Pinzon, W. L. Roles of Cytokines in the Pathogenesis and Therapy of Type 1 Diabetes. Cell Biochem. Biophys. 2007, 48 (2–3), 159–163. 10.1007/s12013-007-0029-2.

(43) Li, D. Q.; Shang, T. Y.; Kim, H. S.; Solomon, A.; Lokeshwar, B. L.; Pflugfelder, S. C. Regulated Expression of Collagenases MMP-1, -8, and -13 and Stromelysins MMP-3, -10, and -11 by Human Corneal Epithelial Cells. Investig. Ophthalmol. Vis. Sci. 2003, 44 (7), 2928–2936. 10.1167/iovs.02-0874.

