## Supplemental Figures for "Stressed β-cells contribute to loss of peri-islet extracellular matrix in type 1 diabetes"

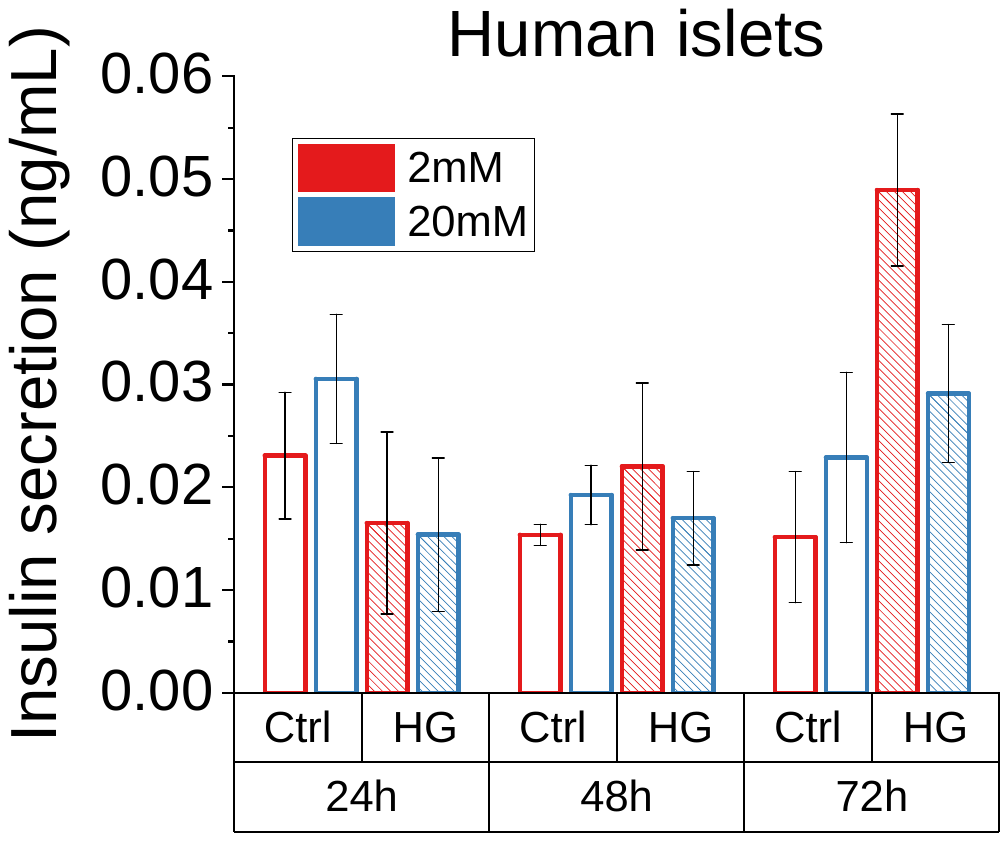

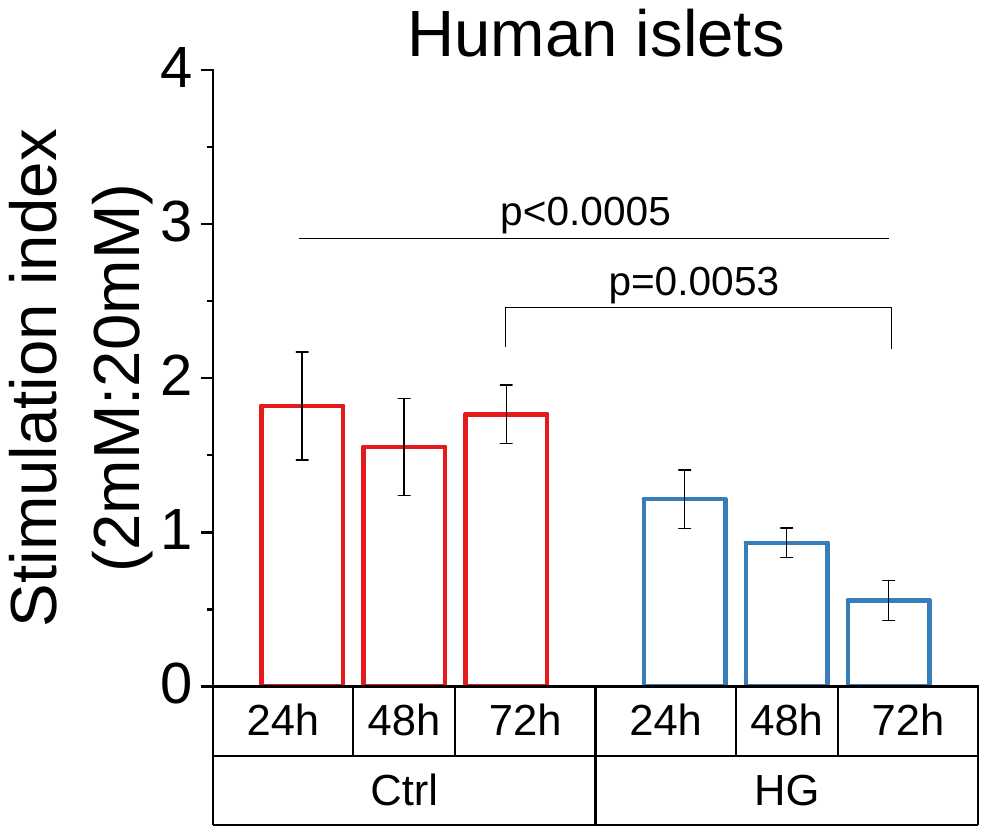

**A**

**B**

**Supplemental Figure 1:** (A) Secreted insulin normalized to insulin content for control and hyperglycemic human islets treated with excess glucose at 24, 48, or 72h at non-stimulatory (2 mM) and stimulatory (20 mM) glucose concentrations (n=3). (B) The stimulation index, or the ratio of insulin secreted at 20 mM glucose to 2 mM glucose, of the same treatment groups in A (n=3). Error bars represent the mean +/- SEM.

**Insulin**

**COL IV**

**Merged**

**Nuclei**

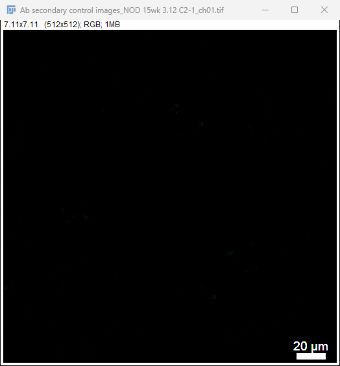

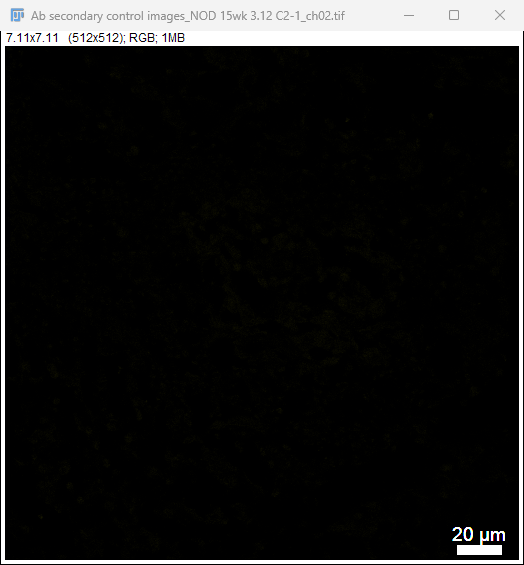

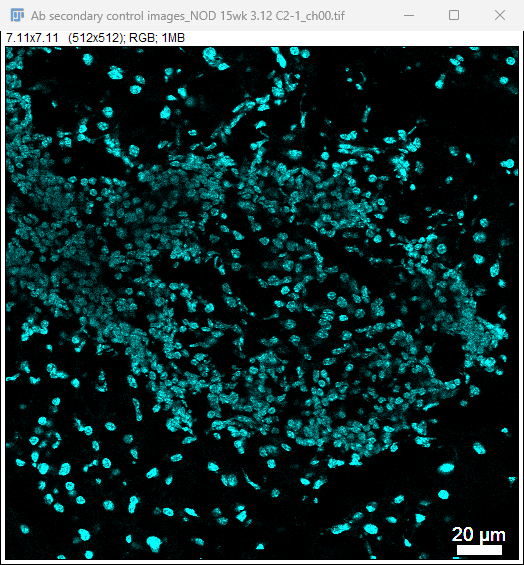

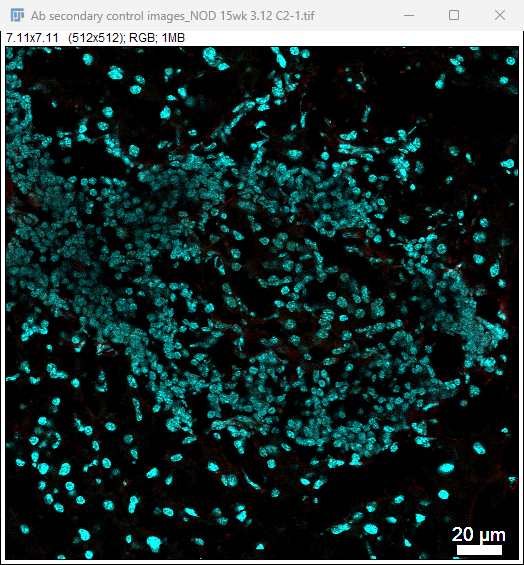

**Nuclei**

**Insulin**

**MMP-3**

**Merged**

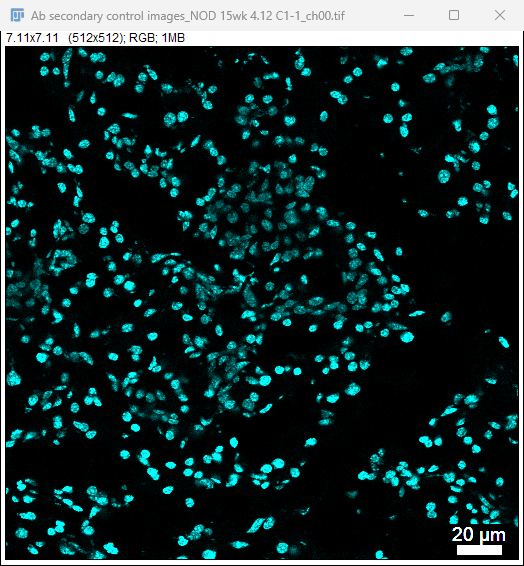

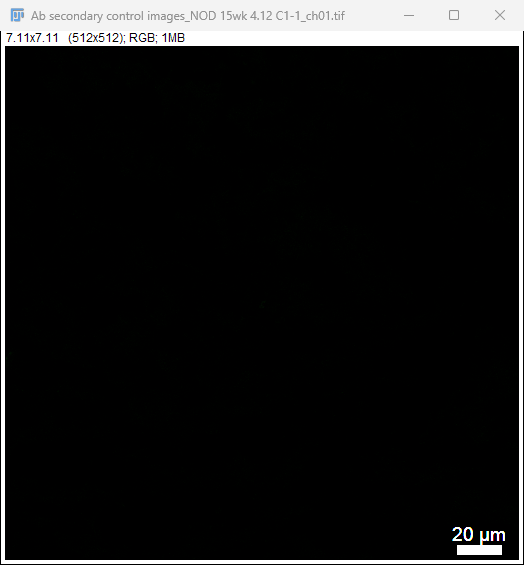

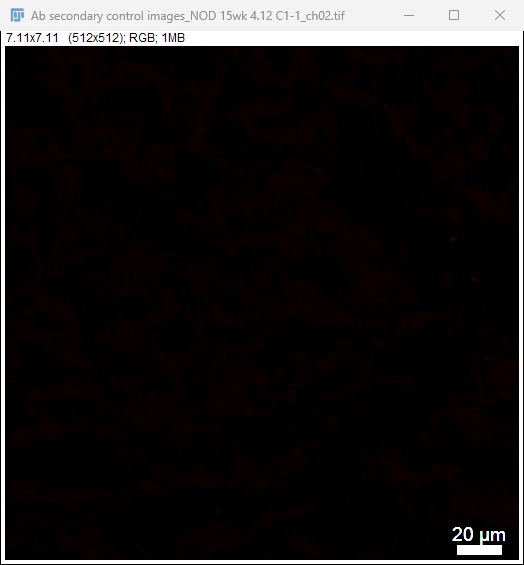

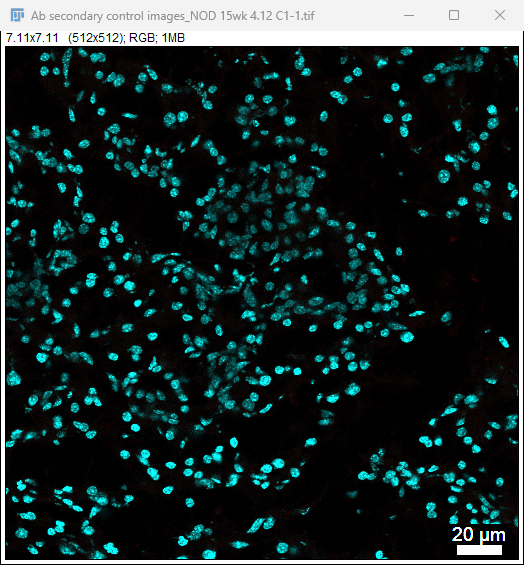

**A**

**B**

**Supplemental Figure 2:** (A) MMP-3 and (B) COL IV antibody controls in mouse sections for immunohistochemistry. All scale bars are 20μm.

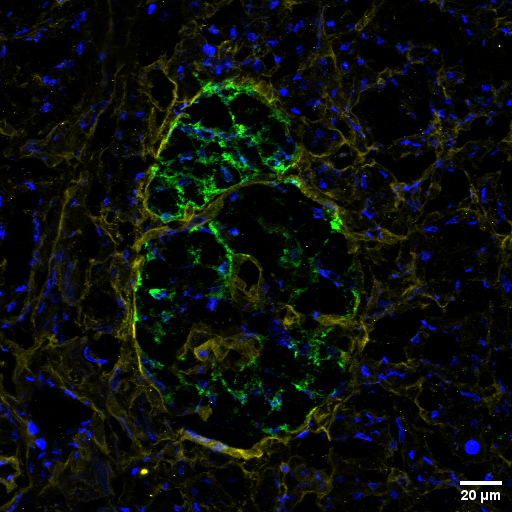

**Ins/COL IV/DAPI**

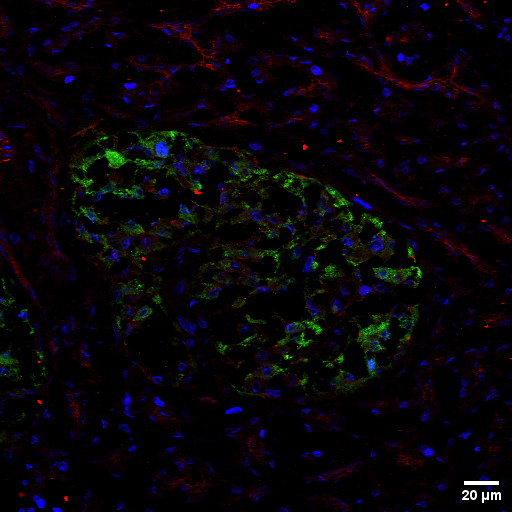

**Ins/MMP-3/DAPI**

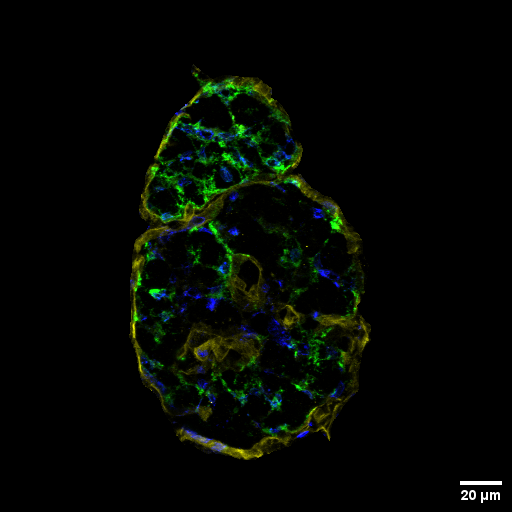

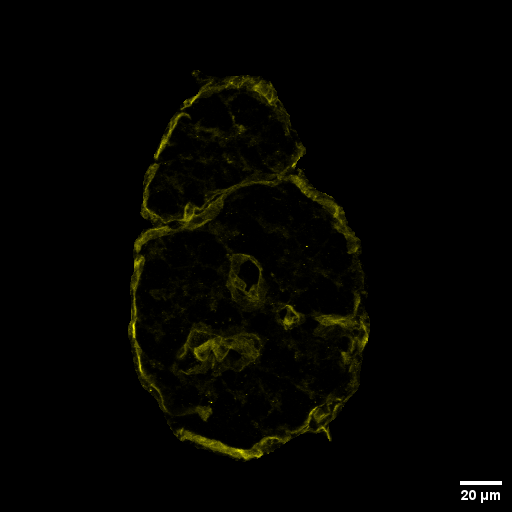

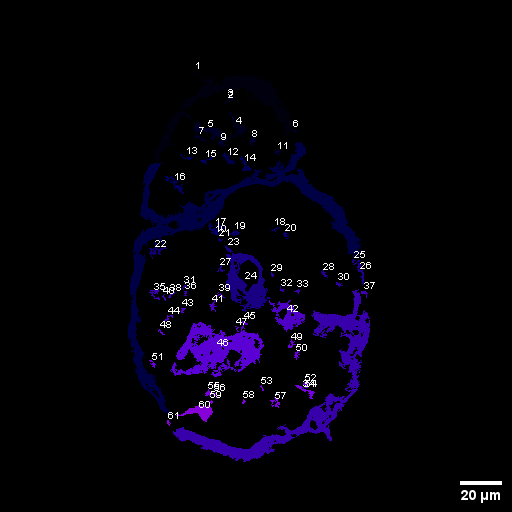

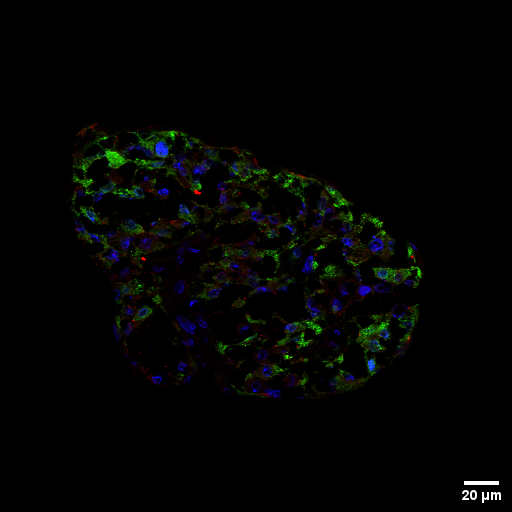

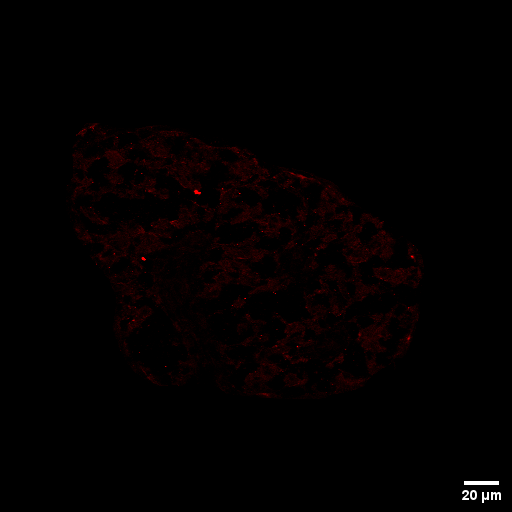

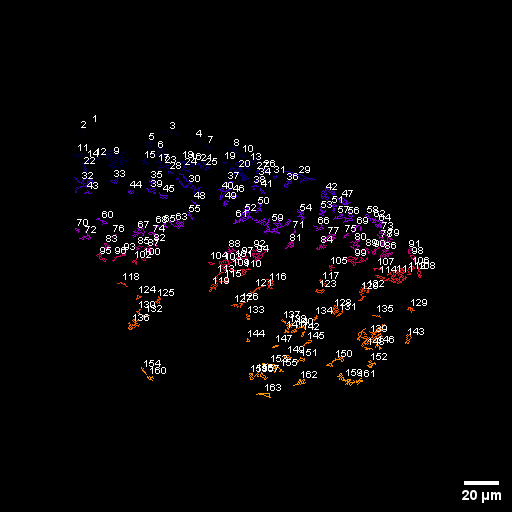

**A**

**B**

**C**

**D**

**E**

**F**

**G**

**H**

**Supplemental Figure 3:** (A) Representative image of insulin (green), MMP-3 (red), and nuclear (DAPI, blue) staining of human pancreas sections. (B) Image in A after removal of fluorescence signal outside of insulin positive islet area. (C) Isolated MMP-3 fluorescence from the image in B. (D) Identification of MMP-3 positive areas within the islet from the image in C using ImageJ 3D Objects Counter. (E) Representative image of insulin (green), COL IV (yellow), and nuclear (DAPI, blue) staining of human pancreas sections. (B) Image in A after removal of fluorescence signal outside of peri-islet ECM capsule as determined by insulin signaling and capsule morphology. (C) Isolated COL IV fluorescence from the image in B. (D) Identification of COLIV positive areas within the islet from the image in C using ImageJ 3D Objects Counter. All scale bars are 20μm.

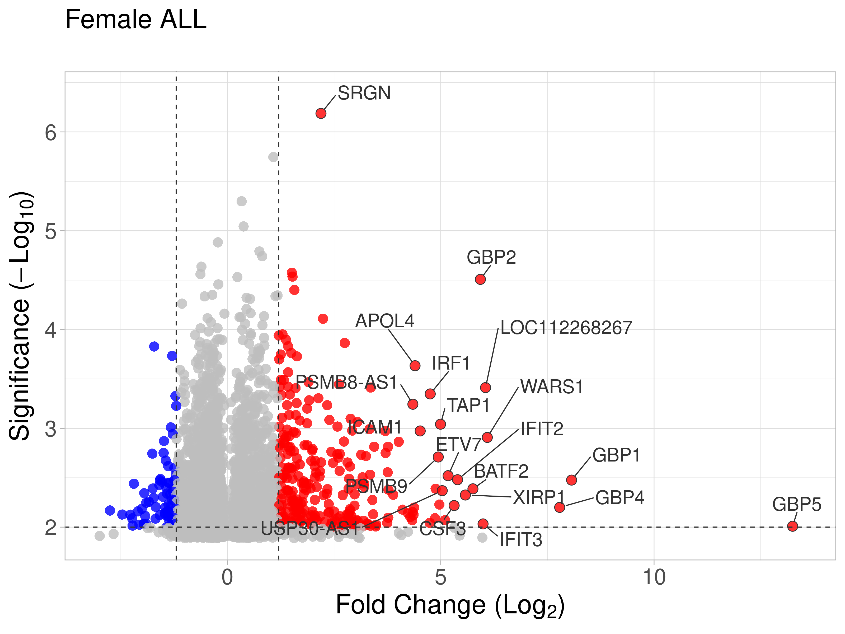

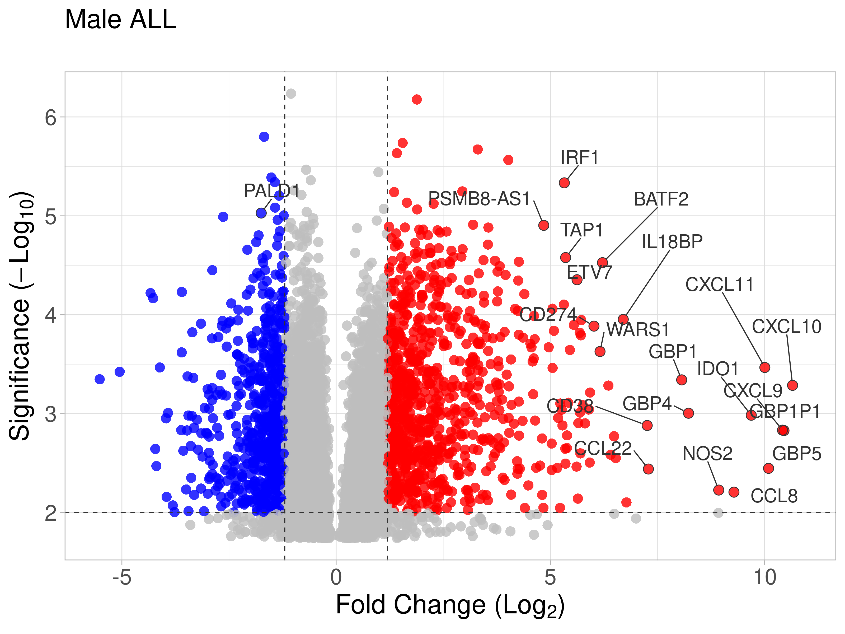

Male

Female

**Supplemental Figure 4:** Volcano plots of all statistically significant transcripts in cytokine treated islets compared to untreated controls from (A) female donors and (B) male donors. Transcripts represented by a red dot are upregulated and blue dots are downregulated upon cytokine treatment in human islets.

**B**

**A**
